# High-resolution microbial network analysis defines biocontrol consortia in the wheat phyllosphere

**DOI:** 10.1101/2025.09.05.674409

**Authors:** Luzia Stalder, Anna Spescha, Monika Maurhofer, Daniel Croll

**Author notes:** Netherlands Institute of Ecology (NIOO-KNAW), Wageningen, the Netherlands.

## Abstract

Plant-associated microbiomes comprise diverse microbial species that coexist and interact, influencing community structure and host plant health. However, our understanding of these interactions in field conditions and at strain-resolution remains limited. This hinders the development of effective biocontrol applications, as laboratory findings often fail to translate to field settings due to insufficient insights into *in situ* interaction network structures. This study addresses this limitation by employing taxon-specific high-resolution amplicons to resolve a cross-kingdom co-occurrence network within the wheat phyllosphere microbiome. We performed in-depth monitoring of strains from the hub genus *Pseudomonas*, revealing a high degree of strain-specificity of *Pseudomonas* interactions both within and across kingdoms. Through negative interaction modelling, we identified a consortium of ten biocontrol taxa with the potential to suppress seven fungal pathogens in field conditions. Additional stabilizer strains were found to likely enhance persistence. We validated the strain-specific interactions of *Pseudomonas* with the major fungal pathogen *Zymoseptoria tritici* using co-inoculation experiments with genotypes retrieved from the same field. Consistent with our prediction, we identified a *P. poae* isolate as the most antagonistic towards the pathogen both *in vitro* and *in planta*. Our study demonstrates that taxon-specific high-resolution network inference can effectively map microbial interaction networks and predict strain-specific interaction patterns of biocontrol genotypes with high persistence under field conditions. Our novel approach supports the design of more effective and sustainable biocontrol strategies.

## Introduction

Plants are inhabited by a diverse consortium of microorganisms, including bacteria and fungi, which together form complex interaction networks. Interactions within these ecological networks can have varying impacts on the species involved, ranging from positive over neutral to negative[1]. For instance, mutualism, a beneficial relationship for both parties, occurs when two species exchange metabolic products for mutual benefit or cooperate to construct a biofilm that provides its members with antibiotic resistance[2, 3]. Commensal relationships, where one partner benefits without affecting the other, are often observed in biodegradation processes, where commensals feed on compounds produced by other community members[4]. Competition occurs when two species with similar niches exclude each other, which can be mediated through various mechanisms[5]. Plant-associated bacteria can engage in direct antagonistic interactions mediated by contact-dependent killing mechanisms, largely mediated by the bacterial type VI secretion system, a molecular weapon deployed by many Proteobacteria to deliver toxins into both eukaryotic and prokaryotic cells[6]. Moreover, numerous plant-associated microbes have been shown to secrete chemical compounds that directly suppress the growth of microbial opponents[5]. In addition to antibiotic production, different bacteria including *Pseudomonas* and *Streptomyces* can also produce volatile organic compounds (VOCs) that act as information signal within and between microbial groups and have been shown to inhibit the growth of a broad diversity of plant-associated fungi[7]. Microbes can also use indirect mechanisms to compete with other microbes, such as rapid and efficient utilization of limited resources. For instance, bacteria have developed complex methods to capture iron by releasing siderophores, which in turn modifies the growth of competing microbes in their close surroundings[8, 9]. For example, the secretion of iron-chelating molecules by beneficial *Pseudomonas* species has been linked to the suppression of diseases caused by fungal pathogens[10]. Most of these mechanistic insights on microbial interactions stem from controlled laboratory experiments conducted in reduced systems with few model species and hosts. However, it is only poorly understood in which contexts these interactions take place in actual ecosystems [11, 12]. Moreover, the dynamics and stability of interactions in ecosystems is largely unexplored.

The crop microbiome has been a plant ecosystem of major interest due to its relevance for food security[13, 14]. During the process of domestication, both the crop plants and their associated microbiomes experienced a significant reduction in diversity[15]. This decrease is associated with a change in species interactions and specifically a higher likelihood of species invasion and pathogen infestation[16]. In cereals, fungal pathogens pose a major concern[17]. For instance, in wheat, the fungal pathogen *Zymoseptoria tritici* is one of the most dominant fungi[18–20]. *Z. tritici* is the most damaging fungal wheat pathogen in Europe and is also a significant wheat pathogen worldwide[21, 22]. The rapid emergence of fungicide resistance has necessitated alternative control strategies. Hence, understanding its interactions with other microbiome members, especially potential antagonists, could open new avenues for disease control[23]. Biocontrol describes the use of natural antagonists – biological control agents - to manage disease[23, 24]. In wheat, the bacterial genus *Pseudomonas* is particularly dominant[25, 26]. Many *Pseudomonas* species are well-known biocontrol agents due to their ability to suppress plant pathogens[24, 27, 28]. The majority of these suppressive strains are phylogenetically classified within the *P. fluorescens* and *P. putida* groups. A prominent example is *P. chlororaphis*, a member of the *P. fluorescens* group, renowned for its production of a variety of phenazines and volatile organic compounds (VOCs) that directly inhibit microbial growth[29–32]. However, the interactions of *Pseudomonas* with microbiome members in the field, particularly fungi, are not well understood[33, 34]. While previous studies suggest that these interactions could be species and even strain-specific, systematic field studies to confirm this hypothesis are lacking[35, 36]. Moreover, many attempts to translate biocontrol interactions from laboratory experiments to the field have not been successful, strongly suggesting that interactions are dependent on the surrounding microbiome[37, 38].

So far, complex microbial interactions in the field have been mostly studied based on co-occurrence[12, 39]. The basic principle of co-occurrence modelling is to identify patterns of species that tend to occur together more or less frequently than would be expected by chance. Importantly, both positive and negative interactions can result from different underlying causes. Co-occurrence can arise through direct interactions such as competition for resources, predation or mutualism, but also from indirect interactions including amenalism or commensalism[12]. Co-occurrence of two microbes can also arise from simply sharing an environmental niche, hence the correction for environmental factors is essential for meaningful interpretation of co-occurrence patterns[12, 40]. Moreover, the sampling scale critically determines co-occurrence patterns that can be observed. It has been argued that interactions acting at the individual scale as a local process might not be discernible at coarser spatial scales[41–43]. Hence, resolution needs to be carefully chosen so that the desired co-occurrence signal can be extracted from the data.

The most common approach to measure co-occurrence patterns in the field is amplicon sequencing of ribosomal markers[44]. Amplicon sequencing allows for abundance monitoring of microbes directly in environmental samples. Despite its advantages, amplicon sequencing based on either 16S ribosomal DNA for prokaryotes or internal transcribed spacers (ITS) for eukaryotes is often limited by its inability to resolve below the genus level[44, 45]. Metagenomic approaches can offer a higher resolution of microbial communities if the targeted organisms are abundant, but they too lack species resolution, as the assemblies represent a consensus sequence of many similar taxa[44]. This limited resolution poses a significant problem as microbial interactions are often species or even strain-specific[35, 36].

In this study, we address this issue by employing pangenome-informed taxon-specific amplicon sequencing[46]. This innovative approach enabled us to achieve species and strain resolution of major taxa from field samples. By combining this method with co-occurrence modelling, we infer the species-specific cross-kingdom interaction network within the wheat phyllosphere microbiome, focussing on the major fungal pathogen *Z. tritici* along with the dominant *Pseudomonas* genus. Furthermore, we sought to identify persistent biocontrol genotypes and stabilizer taxa that exhibit positive interactions with these candidates. We validate the predicted strain-specific interactions of *Pseudomonas* with *Z. tritici* using co-inoculation experiments with genotypes retrieved from the same field. We show that utilizing a comprehensive isolate collection from the same field helps to experimentally validate strain-specific interactions identified through high-resolution network analyses of amplicon sequencing data.

## Methods

### Wheat field sampling scheme

We analyzed eight elite European winter wheat (*Triticum aestivum*) varieties sampled at five different timepoints over the growing season as described previously[46]. Specifically, the wheat was sampled on the 23.05.2019 (growth stage Feekes 7.0), 06.06.2019 (Feekes 10.0), 27.06.2019 (Feekes 10.5), 15.07.2019 (Feekes 11.2) and 22.07.2019 (Feekes 11.4)[47]. The sampled cultivars included Aubusson, Arobase, Lorenzo, CH Nara, Zinal, Simano, Forel and Titlis. Two biological replicates of the wheat panel were grown in two complete block designs separated by approximately 100m at the field phenotyping platform site of the Eschikon Field Station of the ETH Zurich, Switzerland (coordinates 47.449°N, 8.682°E). The cultivars were grown in plots of 1.2-by-1.7-m, with the genotypes arranged randomly within each block. No fungicides were applied. For each cultivar, block and timepoint, two plants were collected. From each plant three leaves were collected: the bottom leaf touching the ground (first leaf), the lowest leaf not touching the ground (second leaf) and the flag leaf (fourth leaf). Each leaf was immediately stored in a plastic foil to avoid contamination, stored at 4°C overnight before processing. In total, 480 leaves were collected (8 cultivars x 2 blocks x 5 timepoints x 2 plants x 3 leaves). Leaves were fixed on A4 paper with printed reference marks, covered by a transparent foil and scanned with a Canon CanoScan LiDE220 scanner set to a picture resolution of 1200 dpi. The leaf-toolkit v.0.2.0 software was used to analyze the number of pycnidia and the area covered by lesions[48]. Samples were transported on ice, stored at 4°C over night and processed the next day.

### Sample homogenization, DNA extraction, amplification, pooling and cleanup

Leaves were processed as described previously[46]. Leaves were lyophilized for 48 h and weighed. Then, the complete leaves were homogenized using 0.5 mm and 0.2 mm zirconium beads in the bead ruptor bead mill homogenizer (OMNI) using the following settings: speed 5.00, number of cycles 2, time of cycle 1:00, time distance between cycles 1:00. DNA extraction was performed with automated magnetic-particle processing using the KingFisher Flex Purification Systems (Thermo Scientific). To enhance the DNA extraction of fungal and bacterial DNA, lyticase and lysozyme was added to the first lysis step with PVP buffer. Specifically, for 10 mg dry leaf mass 3.9 µl lyticase (200,000 U/mg, diluted to 6.5 mg/ml), 3.9 µl lysozyme (22,500 U/mg, diluted to 10 mg/ml) and 98 µl PVP lysis buffer was added, and samples were incubated at 55°C for 30 min. Then, 3.9 µl proteinase K (30 U/mg, diluted to 10 mg/ml) for 10 mg dry mass was added and incubated at 55°C for 30 min. From each sample, 150 µl of clear lysate was transferred to an empty binding plate (KingFisher Flex, Thermo Scientific). For each sample, 360 µl PN binding buffer, 30 µl well suspended Sbeadex beads were added. Using the KingFisher Flex, the first washing step was performed using 400 µl PN1 buffer per sample, then a second wash using 390 µl buffer PN1 with 10 µl RNase A (diluted to 10mg/µl in nuclease-free water) and a third wash using 400 µl PN2 buffer. Each sample was eluted in 100 µl nuclease-free water. The DNA concentration was measured using the Spark Microplate reader (Tecan). Then, DNA concentrations were diluted to 5 ng/µl using the Liquid Handling Station (BRAND). PCR reactions were pipetted using the Mosquito HV liquid handling robot (SPT Labtech). The first amplicon PCR reaction was performed in a 15 µl reaction volume. Specifically, 7.5 µl KAPA HiFi HotStart ReadyMix (2x), 1.5 µl forward primer (3 µM), 1.5 µl reverse primer (3 µM), 3 µl DNA (5 ng/µl) and 1.5 µl HPLC water were combined. Primer sequences and cycling protocols are documented in Supplementary Table 1 and 2. All primers were synthesized by IDT (Integrated DNA Technologies, Coralville, IA). *Pseudomonas* and *Z. tritici*-specific amplicon PCR products were diluted 1:5, PCR products from 16S and ITS 1:10. The second barcoding PCR reaction was performed in a 25 µl reaction volume. Specifically, 12.5 µl KAPA HiFi HotStart ReadyMix (2x), 2.5 µl M13 forward barcode (3 µM), 1.5 µl M13 reverse barcode (3 µM), 2 µl diluted PCR product and 5.5 µl HPLC water were combined. The sample meta data with the associated barcode sequences are available in Supplementary Table 3 and 4. Positive and negative extraction controls as well as PCR controls were included on every plate. Samples were pooled by amplicon taking 1.5 µl from each barcoded product. Each amplicon pool was cleaned using AMPure XP beads using a bead ratio of 0.8x. For the first run, the target specific amplicons and the 16S and ITS amplicons were combined in a ratio 3:2. For the second run, only the *Pseudomonas*-specific *rpoD* amplicon and the *Z. tritici* specific amplicon chr. 13 were sequenced in equimolar amounts to increase sequencing depth.

### Library preparation and sequencing

Library preparation and sequencing was carried out at the Functional Genomics Centre Zurich (FGCZ). Two SMRTbell libraries were prepared for each amplicon length using the SMRTbell prep kit 3.0. One for the long 3-kb amplicons (*Pseudomonas*-specific *rpoD* amplicon and *Z. tritici*-specific amplicon on chromosome 13), and one for the 1.5kb 16S and ITS amplicons. Size selection was performed using BluePippin (Sage Science) with a 0.75% dye-free cassette for each library. PacBio sequencing was performed on a Sequel II machine with the SMRT 8M cell. Two sequencing runs were performed, the first one using SMRT Link version 10.1 and the second one SMRT Link version 11.1.

### PacBio raw read processing

CCS were extracted from raw reads using the ccs software from the bioconda package pbccs provided by the manufacturer (Pacific Biosciences). For the first run pbccs v. 6.0.0 was used and for the second run v. 6.3.0. As the second run was run with the software pbccs v. 6.3.0, it was able to resolve heteroduplex reads in double-stranded and single-stranded Zero-Mode Waveguide (ZMW). To make it comparable to the first run that reported only one sequence per heteroduplex, we also included only one sequence per ZMW for single-stranded reads for all further analyses. We split the CCS reads by barcodes using the software lima 2.0.0 (Pacific Biosciences) with the following parameters lima –log-level INFO –per-read –min-passes 0 –split-bam-named –ccs –different - A 1 - B 3 - D 2 - I 2 - X 0. We assigned the reads to the respective amplicons using BLASTn assignments to reference amplicon from the *P. protegens* CHA0 and *Z. tritici* isolate 1E4[49]. All six reference amplicons were blasted against each read and reads were then assigned to the reference hit with the lowest e-value, the highest length and the highest identity. We removed primer sequences using cutadapt v. 3.4 with the following parameters: cutadapt - a FORWARD_PRIMER_SEQ…REVERSE_PRIMER_SEQ –discard-untrimmed –revcomp[50]. Primers were treated as linked, i.e. reads without primers at both ends were discarded.

The R package dada2 v. 1.3.0 was used to infer ASVs for each amplicon separately[51]. In the following, dada2 steps for each amplicon are described. Reads were filtered and trimmed using the function filterAndTrim with the parameters minLen=minLength, maxLen=maxLength, rm.phix=FALSE, maxEE=2, qualityType = “FastqQuality”, multithread=TRUE. Length ranges for each amplicon are described in Supplementary Table 5. Reads were dereplicated using the function derepFastq with the parameters verbose=TRUE, qualityType=“FastqQuality”. Error models were estimated with the function learnErrors with the parameters errorEstimationFunction=dada2:::PacBioErrfun, BAND_SIZE=32, multithread=TRUE. Reads were denoised using the function dada with the parameters BAND_SIZE=32, multithread=TRUE, pool=FALSE. ASV sequence tables were generated using the function makeSequenceTable. To remove chimeras from the sequence table, the function removeBimeraDenovo with parameters method=“consensus”, minFoldParentOverAbundance=3.5, multithread=TRUE, verbose=TRUE was used. Total read numbers passing all quality filtering are indicated in Supplementary Table 6. The sequencing run had no significant impact on abundance (Kruskal Wallis p-values n.s.).

### Taxonomic classification

We assigned 16S reads using the dada2-formatted Silva database v. 138[52] and ITS reads to the UNITE database v. 8.3[53]. To perform taxonomy assignment, we used the function assignTaxonomy from the dada2 package. As the UNITE database comprises mostly ITS1-ITS2 reference sequences and only few full-length ITS-LSU sequences, we truncated all ITS reads to the ITS1-ITS2 fragment for assignment. We blasted the reads against the ITS1-ITS2 fragment of *Z.tritici* strain S-46 (KT336200.1) and used the hit coordinates to truncate reads with the seqkit software function subseq[54]. We assigned *Pseudomonas* reads of the *rpoD*, transporter and 16S amplicons to *Pseudomonas* species using BLASTn against all 1071 complete *Pseudomonas* genomes available from the *Pseudomonas* db v. 21.1 (Date of download: 19.02.2023, Supplementary Table 7)[49, 55]. The best assignment was chosen according to BLASTn bitscores. *Pseudomonas* species were assigned to groups and subgroups according to Lalucat et al. 2020[56]. We assigned *Z. tritici* reads of the *Z. tritici* amplicons on chromosomes 9 and 13, and of the ITS amplicon to a database of previously sequenced *Z. tritici* strains. For this, we used draft assemblies of previously collected 177 genomes from the Eschikon Field Station of the ETH Zurich, Switzerland[57], as well as genomes from the reference pangenome[58]. The best assignment was chosen according to bitscores. Fungal pathogens were classified as phytopathogens based on FUNGuild annotations [59], with additional manual annotations for genera not covered, as detailed in Supplementary Table 8.

### Isolation and characterization of *Pseudomonas* isolates

Leaves were collected as stated above. All eight cultivars and three canopy heights were sampled at two timepoints, that was on the 19.06.2019 (growth stage Feekes 10.1) and on the 12.07.2019 (Feekes 11.2). Leaves were grinded individually with 10 ml 0.9% NaCl solution and then plated in a serial dilution series up to 10^-7^ on Kinǵs B agar [60] with 100 mg/l cycloheximide, 13 mg/l chloramphenicol and 40 mg/l ampicillin. Plates were incubated in the dark for 3 days at 18°C. From each dilution series, 40 colonies were picked with as diverse phenotypes as possible and regrown on Kinǵs B agar with 100 mg/l cycloheximide, 13 mg/l chloramphenicol and 40 mg/l ampicillin over night at 24°C under shaking. Colonies were picked and transferred into microtiter plates with 200 µl lysogeny broth (LB) [61]. Glycerol stocks were prepared by adding the same volume of 89% glycerol and then stored at - 80°C. For characterization of the isolates, glycerol stocks were grown in microtiter plates with 200 μl LB media overnight at 24°C under shaking. Cultures were centrifuged at max. speed for 10 min using a Heraeus, Eppendorf centrifuge and supernatant was discarded. 3 µl HPLC water was added to resuspend the pellet and a 1:500 dilution was prepared. The product was used for the *Pseudomonas rpoD* PCR1 reaction as described above. The reaction products were cleaned using NucleoFast plate according manufactures instructions (Machery Nagel, Switzerland). Specifically, 80 µl HPLC water was add onto the membrane of each well. 24 µl of the PCR reaction was added subsequently and the plate was spinned down for 10 min at max. speed using a Heraeus, Eppendorf centrifuge. 100 µl HPLC water was added and the plate was spinned down again for 10 min at max. speed. 22 µl HPLC was carefully dispensed on the membrane and the plate was shaken for 10 min at 600 rpm. 12 µl was pipetted into a new PCR plate and heat-sealed to send to Microsynth (Switzerland) for Sanger sequencing. Primers were enclosed in a separate tube. Forward and reverse sequences were merged by adding ten N as a spacer and were then assigned to species using blast against the *Pseudomonas* sequences described above. Merged sequences and species assignments for isolates tested experimentally are available in the Supplementary Table 9. As *Pseudomonas* isolation controls, the previously characterized strains *P. protegens* CHA0[62, 63] and *P. protegens* PF[64] were used.

### *In vitro* inhibition assay

The ST99CH_3D7 and ST99CH_1A5 strains of *Z. tritici* were assessed in co-inoculation with *Pseudomonas* isolates indicated in Supplementary Table 9. As *Pseudomonas* controls, the previously characterized strains *P. protegens* PF[64] and *P. chlororaphis* PCL1391[64, 65] were used. Fungal spores were incubated in 50 ml yeast sucrose broth YSB for 6 days at 18°C under shaking. Liquid cultures were poured through two layers of sterile medical gaze into 50 ml Falcon tubes to remove fungal hyphae. The cultures were then centrifuged in an Allegra X12R Centrifuge at 2700 rpm and 4°C for 10 min. The supernatant of the *Z. tritici* strains was discarded and the pellets dissolved in around 20 ml deionized distilled water (ddH_2_O). The concentration of the spore inoculum was estimated using KOVA Glasstic counting chambers (Hycor Biomedical, Inc., California). 1.5 ml of the fungal suspension with a concentration of 10^8^ spores/ml for *Z. tritici* and was prepared. An overnight LB liquid culture of each *Pseudomonas* isolate was poured into a 50 ml falcon tube and centrifuged for 10 min. at 2700 rpm and 4°C in an Allegra X12R centrifuge and the supernatant was discarded. The pellet was dissolved in 15 ml ddH_2_O, centrifuged again and the supernatant was discarded. The pellet was dissolved in 10 mL ddH_2_O. For each culture, the OD_600_ was measured twice using the spectrophotometer Thermoscientific Genesys 150. 1.5 ml of the final suspension with an OD_600_ of 0.5 ≈ 5x10^8^ cfu/ml was prepared. 100 μl of the fungal spore suspension was evenly spread onto square Potato Dextrose Agar (PDA) plates (39 g/L; BD Difco™, Becton, Dickinson and Company, USA) prepared with ddH₂O using a Drigalski spatula. Afterwards, 5 μl of each bacteria suspension was added to the prepared PDA plate. The plates were incubated until evaluation in the dark with the agar side facing down at 18°C. The plates were photographed six and nine days after inoculation. The pictures were taken with a Canon EOS 90D Kit using the settings: ISO 200, Ap 60, Sh 5.6. The plates were placed on a dark background to make the inhibition zones visible. Inhibition zones were evaluated at three edges with the straight-line tool of Fiji (imageJ2, v. 2.14.0) and normalized against the plate diameter.

### *In planta* co-inoculation assay for STB disease assessment

The ST99CH_3D7 and ST99CH_1A5 strains of *Z. tritici* were assessed in co-inoculation with *Pseudomonas* isolates indicated in Supplementary Table 9. As *Pseudomonas* controls, the previously characterized strains *P. protegens* PF[64] and *P. chlororaphis* PCL1391[64, 65] were used*. In planta* experiments were conducted with the wheat cultivar Drifter. 12 seedlings were grown in each square 11 x 11 x 12 cm plastic pot (Bachmann Plantec AG, Switzerland) containing peat soil (Jiffy soil substrate GO PP7, Netherlands) for 17 days prior to inoculation. 10-day-old plants were fertilized with 2 l of fertilizer solution per 15 pots (2 ml/l, Wuxal Universal-Dünger, Maag-Garden, Switzerland). Growing conditions were the following: 16 h of light, 70% relative humidity, temperature of 18°C during the day and 15°C during the night, and light intensity of 12 kLux. 17 day-old wheat plants were spray-inoculated until run-off with 10 ml inoculation suspensions per pot. The suspensions were one of the following: 10 ml mock suspension (ddH₂O and 0.1% (v/v) Tween-20 (Sigma Aldrich)), 10 ml *Pseudomonas* suspension (5 ml *Pseudomonas* suspension and 5 ml mock suspension), 10 ml *Z. tritici* suspension (5 ml *Z. tritici* suspension and 5 ml mock suspension), 10 ml *Pseudomonas* – *Z. tritici* suspension (5 ml *Z.tritici* and 5 ml *Pseudomonas* suspension). *Z. tritici* suspensions were prepared with a concentration of 10^6^ spores/ml and *Pseudomonas* suspensions were prepared with a concentration of OD_600_ of 0.25 ≈ 2 x 10^8^ cfu/ml, as described above. After inoculation, pots were enclosed within a plastic bag for 72 h to ensure 100% relative humidity. 14 and 21 days after inoculations the second and third leaves were harvested and used for pycnidia and lesion quantification. Per treatment and timepoint, 16 leaves were evaluated. Leaves were fixed on A4 paper with printed reference marks and scanned with a Canon CanoScan LiDE220 scanner set to a picture resolution of 1200 dpi. To analyze the number of pycnidia and the area covered by lesions, the leaf-toolkit v.0.2.0 was used[48].

### Leaf disk co-inoculation assay and qPCR quantification

The *Zymoseptoria tritici* strains ST99CH_3D7 and ST99CH_1A5 were co-inoculated with *Pseudomonas* isolates listed in Supplementary Table 9. *Pseudomonas* controls included the well-characterized strains *P. protegens* PF[64] and *P. chlororaphis* PCL1391[64, 65]. Wheat plants were grown as described above. Second and third leaves of 17-day-old plants were excised, cut into 2 cm segments, and placed on 1% water agar prepared with ddH₂O. Suspensions of *Z. tritici* (1 × 10⁶ spores/ml) and *Pseudomonas* (OD₆₀₀ = 0.25, ∼2 × 10⁸ cfu/ml) were prepared as described above. Each leaf segment received 10 μl of *Z. tritici* suspension or mock (ddH₂O with 0.1% Tween-20, Sigma-Aldrich) evenly spread with a brush and dried under sterile conditions for 5 minutes. Subsequently, 10 μl of *Pseudomonas* suspension or mock was applied similarly and dried for 10 minutes. Plates were incubated in climate chambers at 18°C (day) and 15°C (night) with 16 h light and 70% humidity. Eight leaf segments per treatment were evaluated. After 5 days, leaves were frozen at –80°C prior to DNA extraction.

DNA extraction was performed as described above. The TaqMan gene expression master mix (Nr. 4369016, Applied Biosystems) was used for all qPCR quantifications. Pipet 8 µl mix in an optical plate (Applied Biosystems) as per manufacturer’s instructions and use 2 μl of the DNA extracts per reaction. qPCR primers and probes with cycling protocols are available in the Supplementary Table 10. Primers and probes were ordered from Microsynth (Switzerland). Reactions were measured on the Roche Lightcycler 480 (Switzerland) with the detection format 3 color hydrolysis probe. Color compensation as per manufacturer’s instructions was applied. *Pseudomonas* and *Z. tritici* biomass was first normalized to wheat biomass levels and then further normalized relative to the mock-inoculated control, represented as ΔΔCT. The fold change of co-inoculation versus single inoculation is displayed as ΔΔΔCT:

𝚫𝚫: (PS_leaf1_ – Wheat_leaf1_)

𝚫𝚫𝚫𝚫: (PS_leaf1_ – Wheat_leaf1_) – (PS_leaf mock_ – Wheat _leaf mock_)

𝚫𝚫𝚫𝚫𝚫𝚫: ((Ps_leaf PS1 + Zymo_ – Wheat_leaf PS1 + Zymo_) – (PS_leaf mock_ – Wheat_leaf mock_)) - ((Ps_leaf PS1_ – Wheat_leaf PS1_) – (PS_leaf mock_ – Wheat_leaf mock_))

### Network analysis

Networks were generated using the SpiecEasi software to account for the compositional and zero-inflated nature of microbiome data to reduce false-positive detection rates[66]. The sparse and low rank (SLR) method was implemented to account for and remove the statistical effect of latent variables[66]. In total three networks were calculated, the bacteria-fungal network, the *Pseudomonas*-fungal network and the *Pseudomonas*-*Z. tritici* network (Supplementary Table 8, 11, 12 and 13). For bacteria and fungi, the unnormalized 16S and ITS amplicon data was merged by genus. For *Pseudomonas* and *Z. tritici*, the 200 most abundant ASVs were used for network construction. Cross-domain interaction analysis was performed with the following parameter: nlambda = 50, lambda.min.ratio=1e-2, r=r, lambda.log=TRUE, pulsar threshold = 0.05 and rep number = 30. The networks were created independently, and the b parameter, which determines the impact of latent variables on the covariance matrix, was optimized using iterative network creation with rank 10, 11, 18, 32, 40, 45, 48, 50, 55 and the best model was chosen using the extended Bayesian information criterion (BIC). For the bacterial – fungal network, the network with the lowest BIC (rank 40) yielded no negative interactions, which is why we considered the network with the second lowest BIC (rank 1) for further analysis. This network is less conservative in in the latent factor correction, but still yielded a robust network according to the criteria described below.

To assess network robustness, we calculated the relative Hamming distance between ten replicate inferred networks from subsamples (80% of the samples) and the respective large-sample reference network. The relative Hamming distance was defined as (FP+FN)/P, where FP is the number of false positive edges, FN is the number of false negative edges and P is the number of true edges in the reference network. We found that all networks have relative hamming distances <0.5 (Supplementary Table 14). The structural properties of networks (i.e., degree, modularity, and nestedness) were calculated using the igraph R package v. 2.0.3[67, 68]. Visualizations were performed using igraph and Cytoscape v. 3.10.2[69]

### Alignment and phylogenetic tree construction

We performed multiple sequence alignments using PASTA v.1.9.0 with the following MAFFT arguments -- leavegappyregion -- 6merpair -- maxiterate 0 -- adjustdirection -- reorder and FastTree model-gtr - gamma - fastest [70]. We built phylogenetic trees of Sanger sequences using RAxML-NG[71] with the GTR+G model and 100 bootstrap trees. To construct phylogenetic trees of the PacBio sequences of leaf samples, we used FastTree v.2.0.0[72] with default settings that ensured efficient handling of large datasets.

### Statistics

Chi-squared and Wilcoxon Rank Sum tests tests were performed using the R function chi-squared and kruskal.test from the package stats v. 4.2.2[73]. Shannon diversity was calculated using the function diversity from the R package vegan v. 2.6.4[74]. Permutations were calculated for 1000 iterations using the base R function sample with replace=TRUE[75].

### Visualization

To visualize ASV abundances, the R package phyloseq v. 1.42.0 was used [76]. Counts were normalized by sample using the phyloseq function transform_sample_counts(phyloseq_object, function (x) x / sum(x)). Phylogenetic trees were visualized using the R package ggtree v. 3.6.2 [77, 78]. All other plots were created using the R package ggplot2 v. 3.4.2 [79]. Organism and machine icons were created with BioRender.com.

## Results

### Spatio-temporal wheat microbiome analysis via high-resolution amplicon sequencing

Our objective was to map the antagonistic and synergistic interactions within the wheat microbiome using detailed temporal and spatial profiling. We used pairwise co-occurrences to deduce direct antagonisms and synergisms among specific taxa, potentially revealing new ecological traits. Microbial community niche processes occur at small scales, such as individual plant organs and cultivars, and change over time due to factors like seasonal variations, plant growth stages, and soil condition changes[80–82]. To capture these dynamics, we conducted intensive sampling at a single field site in central Switzerland throughout the growing season (Figure 1A). We collected wheat leaf samples at five key time points, from May (when the first node appears) to July (just before harvest). At each time point, we sampled leaves from three different canopy heights to track the plant’s developmental patterns. High-resolution image scans of the collected leaf samples revealed a lesion symptom variation from 0.1% to 99.9%, and a fluctuation of *Z. tritici* pycnidia per leaf from 0 to 1500 across the sampled timepoints (Figure 1B). Our sampling was carried out across two plot sites, each hosting eight different wheat cultivars in a randomized block design. This approach allowed us to observe the microbiome’s behavior across different wheat varieties and growth conditions.

**Figure 1:**
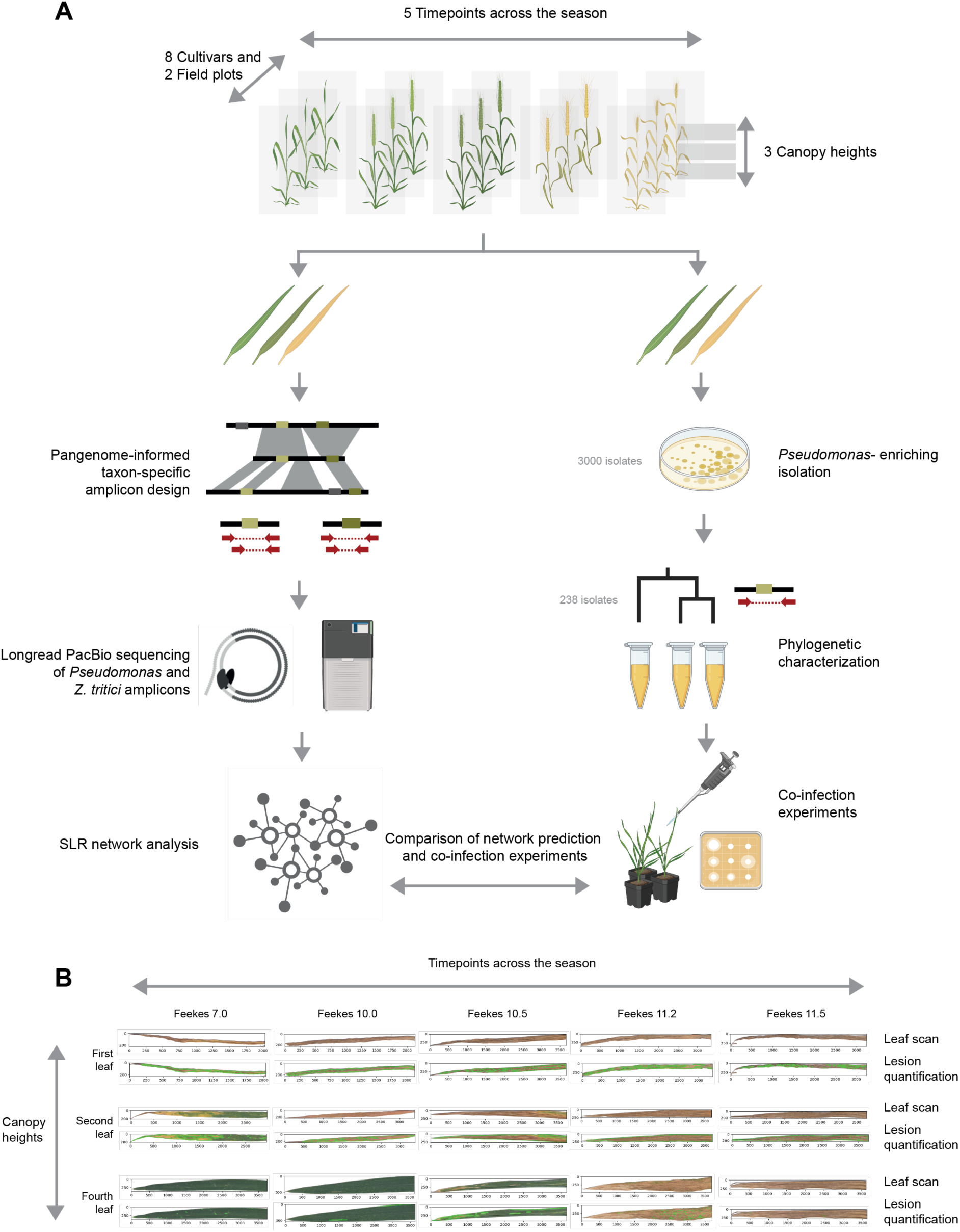
Schematic overview of the study design. (A) Wheat phyllosphere samples were collected at timepoints spanning the main growing season, from May (first node appearance) to July (just before harvest; Feekes growth stages 7.0–11.4). At each timepoint, leaves were sampled from three canopy heights in two field plots, each containing eight wheat cultivars in a randomized block design. Collected leaves were scanned for natural disease symptoms and then used for downstream analyses: either PacBio amplicon sequencing and co-occurrence network analysis (left panel) or *Pseudomonas* isolation for experimental assays (right panel). PacBio sequencing employed pangenome-informed amplicons for *Pseudomonas* and *Zymoseptoria tritici* together with full-length 16S and ITS markers, and networks were inferred with the sparse and low-rank (SLR) method of Kurtz et al. (2019)[40, 66]. For experimental validation, *Pseudomonas*-enriching isolation was performed, isolates were species-assigned with a *Pseudomonas*-specific marker, and strains were tested in both *in vitro* and *in planta* co-inoculation assays with *Z. tritici*. (B) Exemplary leaf samples from the five illustrated timepoints and three canopy heights are shown. Each sample is displayed as the original scan (top) and the corresponding automated lesion detection output (bottom), highlighting *Z. tritici* symptoms (lesion size, pycnidia).

We processed 480 leaves for high-resolution amplicon characterization. We used pangenome-informed taxon-specific amplicon markers to profile the diversity of major bacterial and fungal taxa in high-resolution[46]. We used a *Pseudomonas*-specific amplicon marker around the *rpoD* locus and a *Zymoseptoria tritici*-specific marker on chromosome 13 to resolve diversity within two major bacterial and fungal groups of the wheat microbiome [46]. In addition, we used full-length 16S and ITS ribosomal markers to profile the fungal and bacterial diversity at a general scale. Each reaction was performed in triplicate, resulting in 5,760 amplifications sequenced twice on two PacBio Sequel II flow cells (Supplementary Table 6). This approach enabled us to identify candidates for synergistic and antagonistic interactions at different taxonomic scales. To experimentally characterize potential synergistic and antagonistic interactions discovered at our study site, we established an extensive isolate collection of *Pseudomonas* and *Z. tritici,* sampled within the same season and at the same site where we collected the samples for amplicon sequencing (Supplementary Table 9).

### Bacterial and fungal diversity in the wheat phyllosphere

Our analysis of the wheat phyllosphere revealed a diverse assembly of bacteria. Based on the full-length 16S amplicon, we found *Pseudomonas* and *Sphingomonas* to be the dominant genera (Figure 2A). The *Pseudomonas*-specific amplicon revealed a diverse assembly of *Pseudomonas* species, primarily belonging to the *P. fluorescens* and *P. syringae* groups (Figure 2B and C). The mycobiome, analyzed using the full-length ITS, was predominantly composed of the fungal genera *Zymoseptoria* and *Cladosporium* (Figure 2A). This is broadly consistent with recent findings in wheat fields [19, 20].

**Figure 2:**
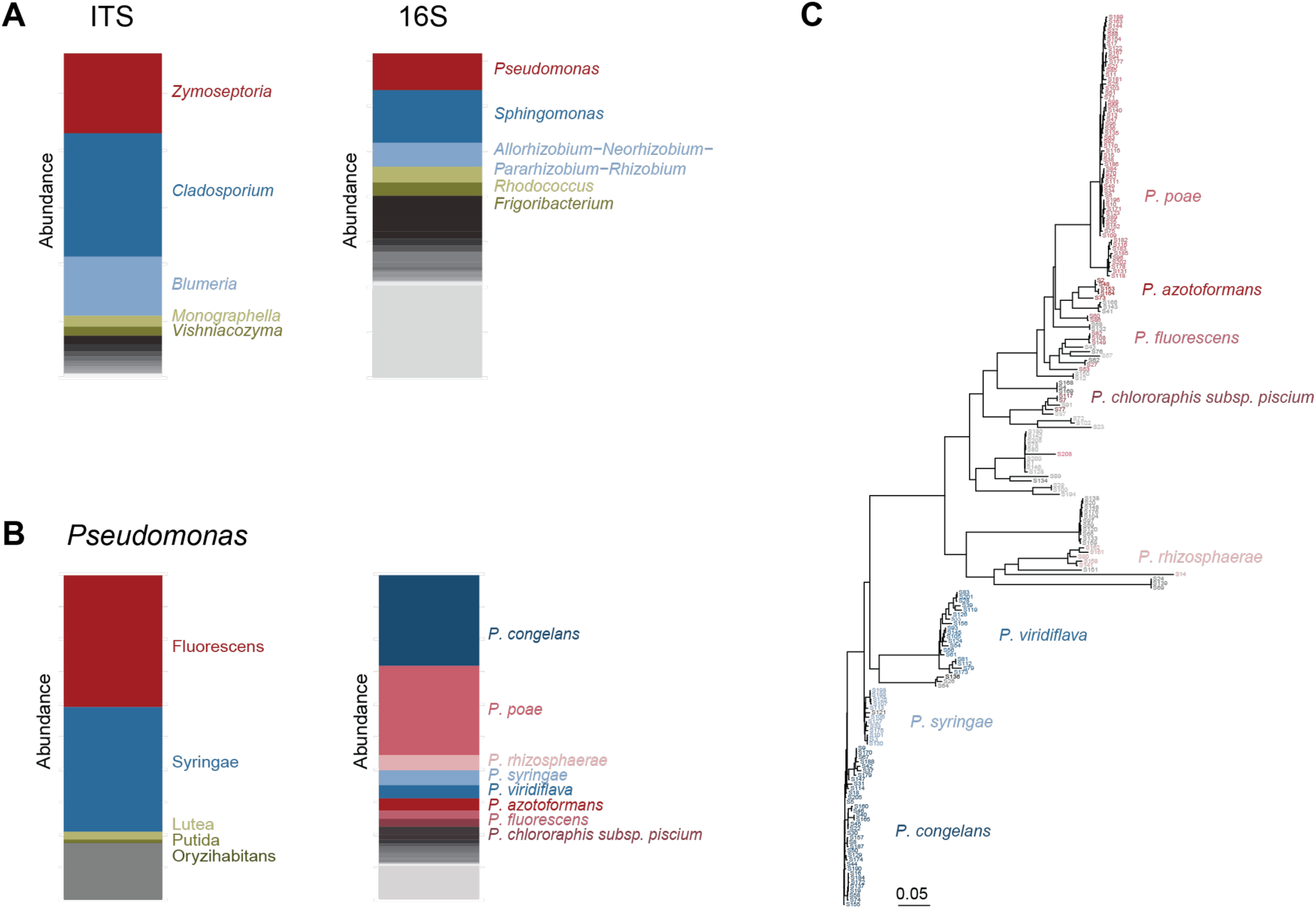
Relative abundances of bacterial and fungal genera in the phyllosphere. (A) Relative abundances of fungal genera determined by the full-length ITS amplicon, respectively bacterial genera determined by the full-length 16S amplicon. (B) Relative abundances of *Pseudomonas* subgroups (left) and species (right) assessed by the *Pseudomonas*-specific amplicon (C) Phylogenetic tree of the 200 most abundant *Pseudomonas* ASVs.

Previous studies have identified a stable phyllosphere core microbiome that remains consistent across various sites and seasons[83, 84]. In accordance with these findings, our analysis revealed no significant variations in the overall bacterial and fungal assemblies when considering different timepoints, cultivars, or canopy heights (kruskal.wallis p-values n.s.). This suggests a stable assembly across season and cultivars. This stability provides us with a reliable baseline for the comparison of specific synergistic or antagonistic co-occurrence patterns.

### Identification of keystone taxa

To determine the extent of synergistic and cross-kingdom interactions between fungi and bacteria on a general scale, we utilized network analysis based on the full-length 16S and ITS amplicon data. Inferring synergistic and antagonistic relationships from amplicon data poses several challenges[40, 66]. First, amplicon-based datasets are compositional and microbial abundances are not independent, hence, traditional correlation metrics for the detection of interactions can lead to spurious results. Secondly, microbial sequencing studies typically measure hundreds of taxa on only tens to hundreds of samples; thus, inference of association networks is severely under-powered. Thirdly, co-occurrence patterns can be influenced by both observed and unobserved environmental factors. To address these complexities, we employed the SpiecEasi Sparse and Low Rank (SLR) method [40]. This method uses a latent graphical model interference scheme that jointly tries to identify robust microbial associations, compositional biases, and technical and environmental covariates influencing microbial associations. Applying this method, we identified 903 significant co-occurrence patters, i.e. putative interactions, among genera (Figure 3A, Supplementary Table 11). We found a dominance of positive interactions (61.9%) over negative interactions (38.1%), and a dominance of within-kingdom interactions (55%) compared to cross-kingdom interactions (45%). The positive interaction network showed higher modularity compared to chance (Supplementary Table 15). Even though it is generally assumed that modularity has a stabilizing effect on network structure, it has been demonstrated that the stabilizing effect is condition dependent[85]. Specifically, in networks with a positive mean interaction strength such as the network found here, a stabilizing effect was not found[85]. The distribution of node degrees showed that most taxa have only a few interactions, while a small number of keystone genera have numerous interacting partners. Among all bacterial genera, *Pseudomonas* exhibited the highest number of interactions (15 interactions). Among the fungal genera, *Blumeria* and *Zymoseptoria* were the most interactive, with 36 and 34 interactions respectively (Figure 3B and 3C).

**Figure 3:**
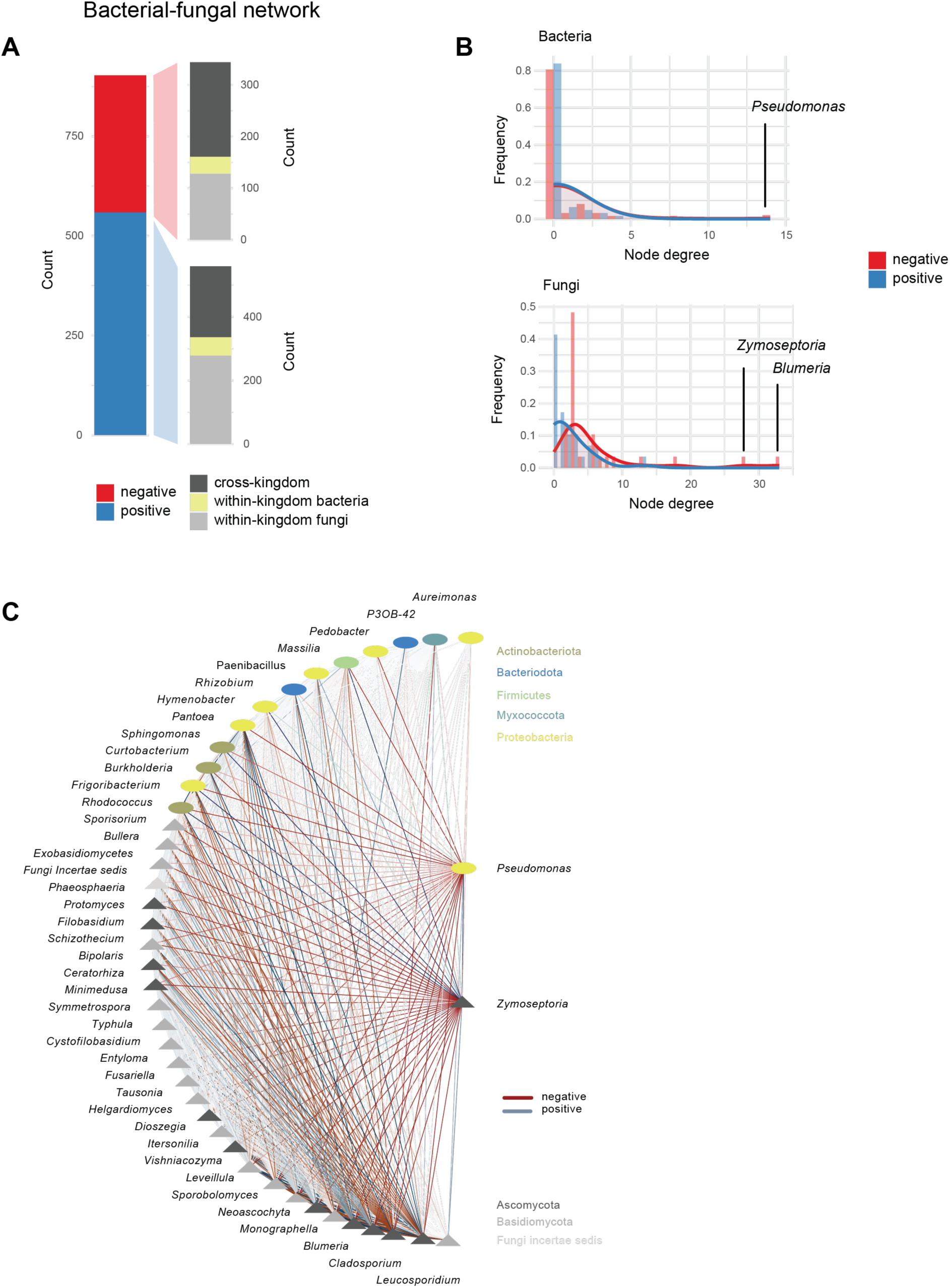
Bacteria-fungal network based on the full-length 16S and ITS amplicon data. (A) Distribution of edge counts within the network. (B) Node degree distribution for both bacterial and fungal genera. The genera acting as hubs, which have the highest node degrees, are highlighted. (C) Interactions of *Zymoseptoria* and *Pseudomonas*. The nodes are ordered based on their kingdom classification and node degree.

### Pathogen suppressor assemblies revealed by *Pseudomonas*-fungal interactions

Next, we wanted to investigate the interactions of *Pseudomonas*, the keystone bacterial genus in the wheat phyllosphere, in more detail. We were particularly interested in whether *Pseudomonas* interactions with fungal genera were specific to certain *Pseudomonas* species or shared among multiple species. To investigate this, we constructed a SLR network of the most abundant *Pseudomonas* Amplicon Sequence Variants (ASVs) detected by the *Pseudomonas*-specific *rpoD* amplicon together with the fungal genera. We included as many of the most abundant *Pseudomonas* ASVs as possible while still ensuring robust network construction (n=200, relative Hamming distance of randomly sampled subnetworks <0.5, see Methods). The resulting network, akin to the previously constructed bacterial-fungal network, was predominantly characterized by positive interactions with a significant modular structure (94.6%), with negative interactions accounting for only 5.4% (Figure 4A, Supplementary Table 12 and 15). Interactions within the *Pseudomonas* genus (80.3%) were more prevalent than *Pseudomonas* cross-kingdom interactions (16.4%). The network topology revealed that *Pseudomonas* cross-kingdom interactions were skewed towards a few taxa with numerous interactions, indicating a species or isolate-specific nature (Figure 4B and C). Conversely, the topology for positive interactions within the *Pseudomonas* genus was less skewed, suggesting a broader range of synergistically interacting species or isolate modules.

**Figure 4:**
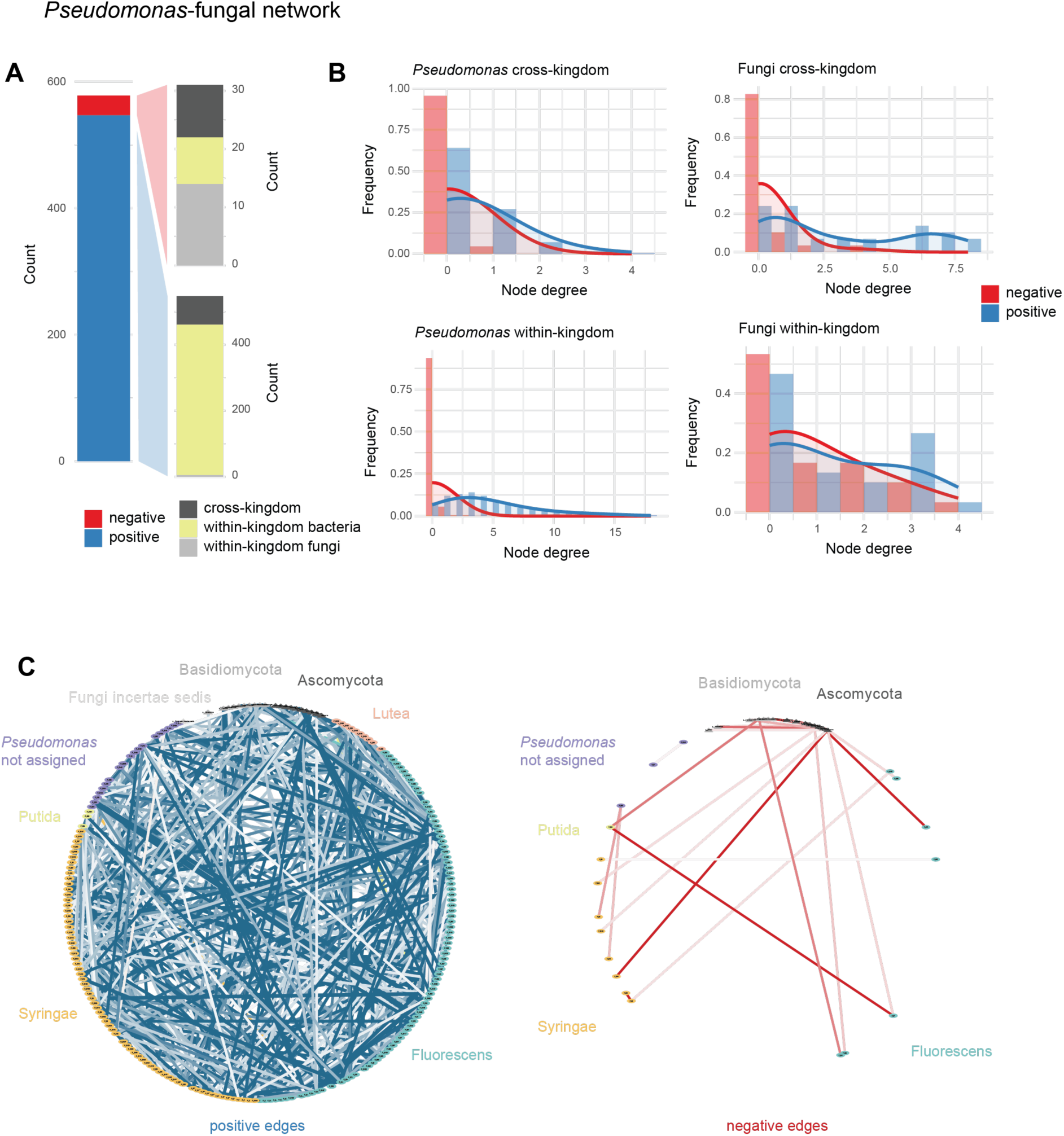
*Pseudomonas*-fungal network based on the *Pseudomonas*-specific amplicon and ITS amplicon data. (A) Distribution of edge counts within the network. (B) Node degree distribution for the *Pseudomonas* and fungal nodes, separated by cross-kingdom and within kingdom interactions. (C) Positive and negative interactions of the network. The nodes are ordered based on *Pseudomonas* group and fungal phyla.

Given the species-specific nature of *Pseudomonas* cross-kingdom interactions, we sought to determine if there was a conserved phylogenetic signal within these interactions. We found that *Pseudomonas* interacting with the same fungal genus are not more closely related than chance, reinforcing the strain-specific nature of these interactions and suggesting a lack of conservation for those interaction patterns among groups (Supplementary Figure 1). Similarly, we found no correlation between phylogenetic distance and interaction weight among *Pseudomonas*-*Pseudomonas* interactions, and similarly no correlation between phylogenetic diversity of *Pseudomonas* ASVs within interaction modules compared to the whole network (Supplementary Figure 1). However, our ability to detect phylogenetic signals of interactions might have been limited due to the correlation of interaction degree and general abundance dominating the phylogenetic signal (Supplementary Figure 2).

Upon examining the negative interactions, we found that most cross-kingdom interactions of *Pseudomonas* were with pathogenic fungal genera (77%, n=7/9, Supplementary Figure 3). Consequently, we systematically assessed pathogen suppressor species and genera (those with only negative interactions with pathogenic fungal genera), and pathogen facilitators (those with only positive interactions with pathogenic fungal genera). We identified a community of ten suppressors or biocontrol candidates, including five *Pseudomonas* ASVs and five fungal genera (Figure 5A and B, Supplementary Table 7 and 12). From the *Pseudomonas* ASVs, three belong to the *P. fluorescens* group (i.e. *P. poae* S70, *P. poae* S95 and *P. chlororaphis subsp. piscium* S7), and two to the *P. syringae* group (i.e. *P. congelans* S44 and *P. congelans* S45). From the fungal genera, four are *Basidiomycota* (i.e. *Sporobolomyce*, *Leucosporidium*, *Filobasidium* and *Bullera*) and one is an *Ascomycota* (i.e. *Schizothecium*). Together, these ten taxa suppress a total of seven pathogenic fungal genera. Furthermore, we identified no negative interactions among the ten suppressors, indicating the potential for a stable sub-community. These suppressors exhibited varying relative abundance patterns across time points, cultivars, and canopy heights (Figure 5C). We also screened the interaction network for suppressor stabilizers, defined as nodes exhibiting exclusively positive associations with suppressors, and identified eight *Pseudomonas* ASVs functioning as suppressor stabilizers, thereby stabilizing six of the ten suppressor taxa (Figure 5B).

**Figure 5:**
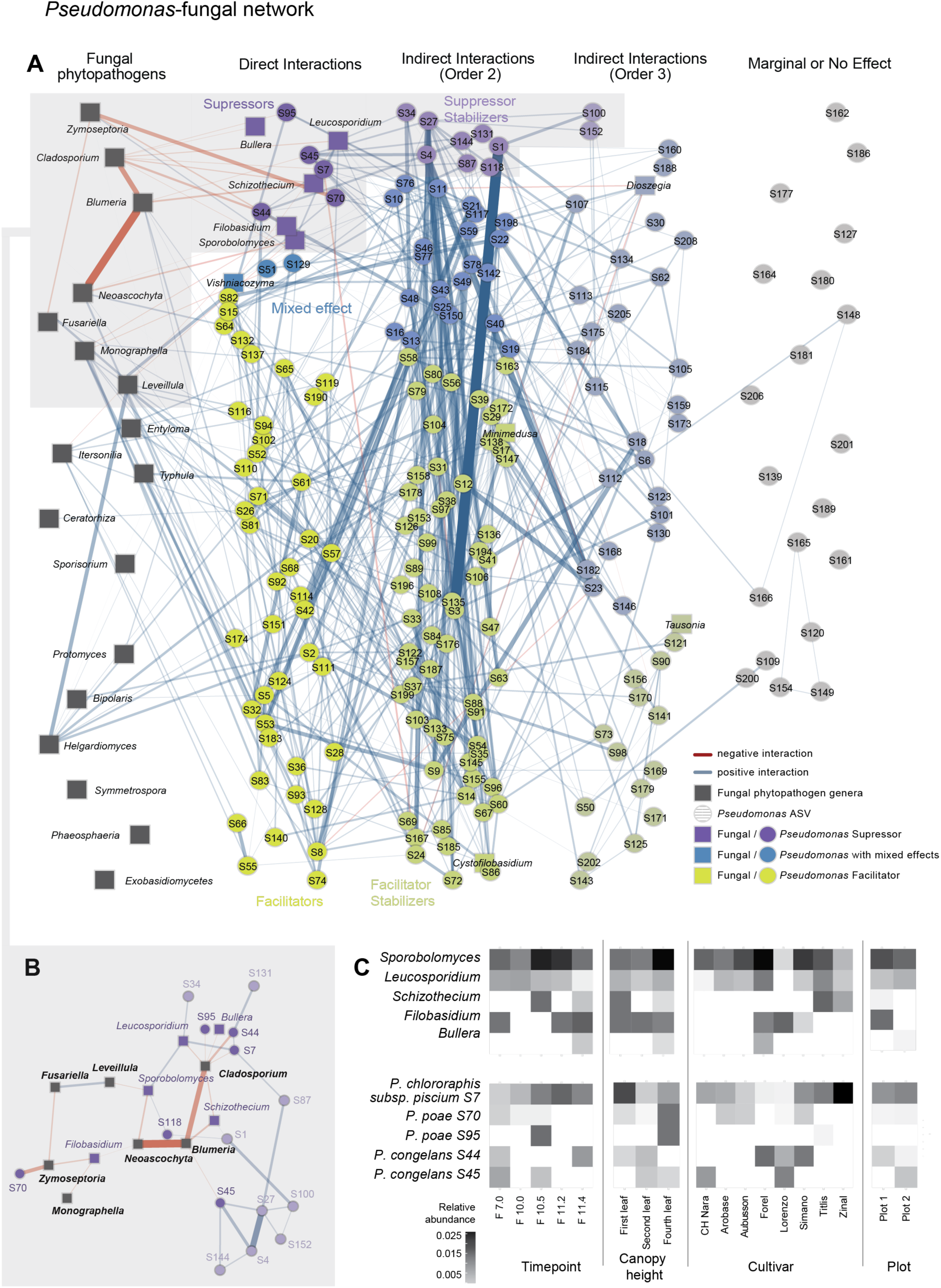
Interactions of fungal phytopathogens. (A) Direct and indirect interactions of genera containing fungal phytopathogenic species with *Pseudomonas* and other fungal genera. Red edges represent negative co-occurrences, while blue edges represent positive ones. The line width is proportional to the strength of the co-occurrence. The first column displays all phytopathogenic fungal genera, represented by dark grey rectangles. The second column shows all direct interactions of these phytopathogenic fungal genera with *Pseudomonas* ASVs (depicted as circles) and other fungal genera (depicted as rectangles). Taxa that only have negative interactions with phytopathogens are labeled as pathogen suppressors and are colored purple. Taxa that only have positive interactions with phytopathogens are labeled as pathogen facilitators and are colored green. Taxa that have both positive and negative interactions with phytopathogens are labeled as having mixed effects and are colored blue. The third column shows the second-order interactions of taxa with taxa that have first-order interactions with phytopathogens. Taxa that have a positive interaction with suppressors are labeled as suppressor stabilizer. Taxa that have a positive interaction with facilitators are labeled as facilitator stabilizer. The same concept applies to third-order interactions. (B) Subnetwork consisting of suppressor and suppressor stabilizer taxa. (C) Relative abundance of suppressor taxa across timepoints, canopy heights, cultivar and plot. Timepoints are indicated according to the Feekes wheat growing stages and canopy heights indicate the bottom leaf touching the ground (first leaf), the lowest leaf not touching the ground (second leaf) and the flag leaf (fourth leaf).

### Cross-kingdom strain-specific interactions

Our previous results have indicated that interactions are often species or strain-specific, leading us to investigate whether we could detect similar level specificity within species. Both positive and negative interactions have been observed between *Pseudomonas* species and *Z. tritici*, a keystone fungal genus in the wheat phyllosphere and a major fungal pathogen[18–20]. To assess *Pseudomonas*-*Z. tritici* interactions in more detail, we constructed a network using the *Pseudomonas*-specific amplicon data as well as the *Z. tritici*-specific amplicon data. *Z. tritici* populations within a single wheat field have shown a pattern of genetic diversity that aligns with a high rate of sexual recombination[57]. Therefore, a sequence variant at the amplicon loci is not associated with other genetic variants in the genome, preventing us from linking traits and functions to individual *Z. tritici* ASVs in our data. However, we hypothesized that the existence of significant interactions would suggest that at least some ASVs represent a biological entity. Our analysis confirmed this, identifying 1634 interactions in total, with 54% occurring within *Pseudomonas* ASVs or within *Z. tritici* ASVs respectively, and 46% being cross-kingdom interactions (Figure 6A, Supplementary Table 13). Similar to the networks constructed at higher taxonomic levels, we found that positive interactions (89.7%) were more common than negative interactions (10.3%). The network topology was skewed towards a few taxa with many interactions, suggesting the existence of keystone taxa that significantly influence network stability (Figure 6B and C). Interestingly, the *Pseudomonas* ASV with the highest number of interactions, identified as *Pseudomonas* sp. 11K1 from the Fluorescens subgroup, interacts positively with 19% of the *Z. tritici* ASVs (38 out of 200). This consistent positive association emphasizes that despite strain-level variability within *Z. tritici*, certain key bacterial taxa can exert a consistently beneficial influence on the pathogen’s community structure.

**Figure 6:**
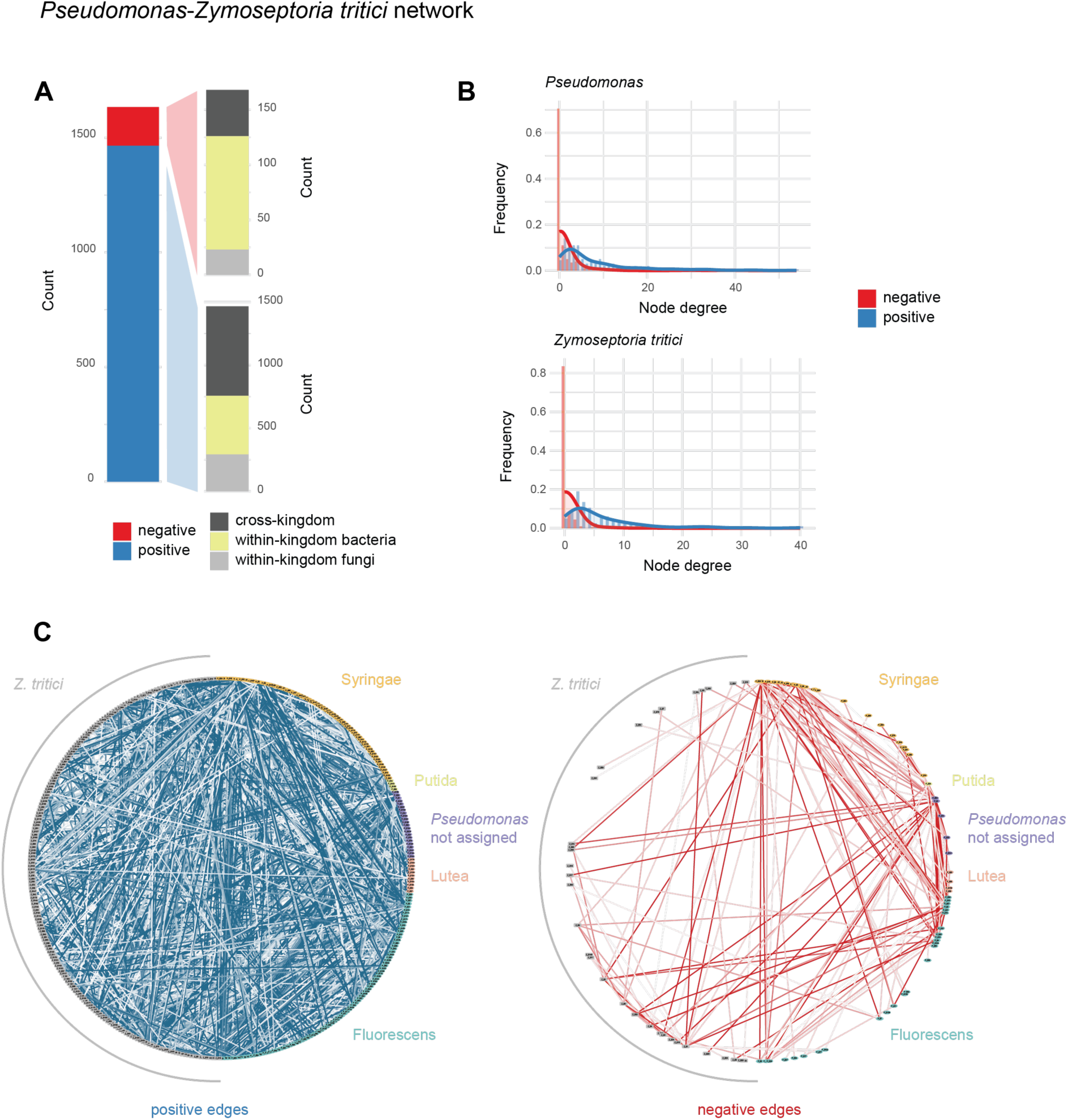
*Pseudomonas*-*Z. tritici* network based on the *Pseudomonas*-specific and the *Z. tritici*-specific amplicon data. (A) Distribution of edge counts within the network. (B) Node degree distribution for *Pseudomonas*, respectively *Z. tritici* ASVs. (C) Positive and negative interactions of the network. The nodes are ordered by *Pseudomonas* group.

### Network analysis predicts experimental co-inoculation outcomes

Our network analysis revealed both species and strain-specific interactions of *Pseudomonas* with different fungal species, including *Z. tritici*. Notably, we found that these strain-specific *Pseudomonas* interactions were also specific to different genotypes of *Z. tritici*. To assess these predictions experimentally, we established a comprehensive collection of *Pseudomonas* isolates from the same field site following the same sampling scheme used for amplicon sequencing. This scheme included sampling at multiple canopy heights and from different wheat cultivars, within the same plots and during the same season as the amplicon data collection. Sanger sequencing of the *Pseudomonas*-specific amplicon marker was then used to assign isolates to species (Figure 7A and B, Supplementary Figure 4 and 5). Although the Sanger amplicon spanned only two-thirds of the PacBio fragment, it was sufficient for reliable species-level identification. Comparison of culture-based and culture-independent amplicon-derived species-abundance distributions revealed significant differences (Figure 7C). In particular, *Pseudomonas congelans* was the most abundant species in the amplicon dataset yet was completely absent from the culture collection of 238 isolates. This discrepancy is likely attributable to primer and amplification bias and/or cultivation bias. A previous evaluation of primer bias, which involved comparing the *Pseudomonas*-specific amplicon around the *rpoD* locus with a second *Pseudomonas*-specific amplicon around an ABC Transporter found it to be non-significant[46]. This suggests that the majority of the discrepancy is likely due to the varying growth abilities of *Pseudomonas* isolates on culture media, a finding that aligns with a study identifying that only a small fraction of *Pseudomonas* species is culturable[86].

**Figure 7:**
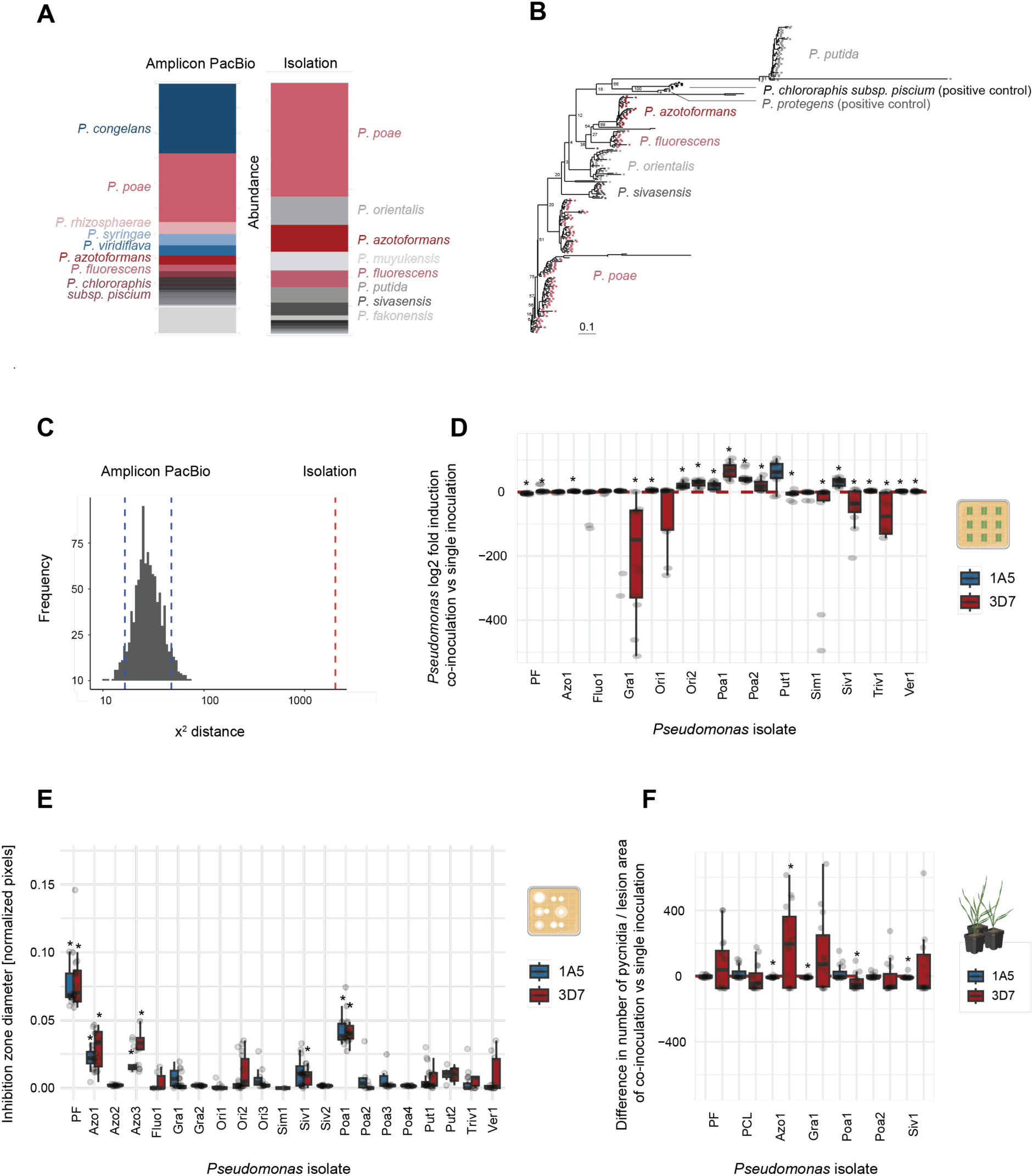
Co-inoculation experiments with *Pseudomonas* spp. and *Zymoseptoria tritici*. (A) Relative abundances of *Pseudomonas* species determined by *Pseudomonas*-specific amplicon sequencing and by isolation frequency. (B) Phylogenetic tree of isolates based on Sanger amplicon sequences. (C) Chi-square distances between the distribution of isolated species and PacBio-based abundance profiles; the null expectation and corresponding 95% confidence interval, derived from random subsampling, are shown. (D) *Pseudomonas* biomass 5 days after co-inoculation with *Z. tritici* strains 1A5 and 3D7, quantified by qPCR and normalized to wheat biomass. Data are shown as log₂ fold induction of co-inoculation relative to single inoculation. (E) *In vitro* growth inhibition of *Z. tritici* strains 1A5 and 3D7 by *Pseudomonas* isolates 5 days post-co-inoculation. Inhibition diameters were measured from plate images and expressed as pixels normalized to plate diameter. (F) Effect of *Pseudomonas* isolates on *Z. tritici* disease severity, measured as pycnidia per lesion area at 21 days post-inoculation. Differences in disease severity (co-inoculation minus single inoculation) are displayed. Isolate acronyms indicate the species group: *P. azotoformans* (Azo), *P. fluorescens* (Fluo), *P. graminis* (Gra), *P. orientalis* (Ori), *P. poae* (Poa), *P. putida* (Put), *P. simiae* (Sim), *P. sivasensis* (Siv), *P. trivialis* (Triv), *P. veronii* (Ver). Isolate characteristics for panels (D)-(F) are given in Supplementary Table 9. *P. protegens* PF served as the bacterial positive control. Statistical significance of co-inoculation versus single inoculation was assessed with Wilcoxon rank-sum tests (p < 0.05).

Out of 238 isolates successfully Sanger-sequenced, 22 were selected to test prediction from our network analysis. To capture the breadth of diversity in our culture collection, we selected representative isolates from each major *Pseudomonas* subgroup recovered in the phylogeny. Within each subgroup, we chose isolates that (i) possessed the longest high-quality Sanger sequences, and (ii) maximized phylogenetic distance from each other to increase within-subgroup diversity. *Pseudomonas* isolates were tested in co-inoculation with two *Z. tritici* isolates from our field study site.

Network analysis identified more positive than negative interactions at both the *Pseudomonas* genus and strain level. To experimentally evaluate the ratio of positive and negative interactions of *Pseudomonas*-*Z. tritici* interactions *in planta*, we co-inoculated *Pseudomonas* and *Z. tritici* on wheat leaf disks and monitored changes in their biomass (Figure 7D and Supplementary Figure 6). Using a triplex Taqman qPCR assay, we simultaneously monitored the biomass of *Pseudomonas*, *Z. tritici*, and wheat (Supplementary Table 10). Consistent with our prediction, we found more positive than negative regulations in co-inoculation (18 positive *vs.* 8 negative median regulations for *Pseudomonas*, and 20 positive vs 6 negative for *Z. tritici*).

Our network analysis identified a *P. poae* isolate as the most antagonistic to *Z. tritici*. We therefore sought to determine whether we could also identify a *P. poae* isolate with antagonistic interactions with *Z. tritici*. We evaluated the antagonism both *in vitro* and *in planta*. For the *in vitro* inhibition, we assessed the inhibition zones of *Z. tritici* caused by different *Pseudomonas* isolates. To test the antagonism against *Z. tritici in planta*, we co-inoculated *Pseudomonas* isolates on wheat leaves to investigate their influence on the development of Septoria Tritici Blotch (STB) disease caused by *Z. tritici*. Consistent with predictions, we found that the most substantial inhibition of *Z. tritici* both *in vitro* and *in planta* was observed by a *P. poae* isolate (Figure 7E-F and Supplementary Figure 7 and 8). Importantly, this inhibition was isolate-specific and not conserved within the species, reflecting the interaction patterns seen in the network analysis.

Our network analysis did not suggest that closely related *Pseudomonas* isolates interact more similarly with fungi than less closely related isolates. Likewise, we tested whether the phylogenetic distance among *Pseudomonas* isolates correlated with a more similar response in our co-inoculation experiments *in vitro* or *in planta*. Consistent with the network analysis prediction, we found no correlation in any of the co-inoculation experiments (Supplementary Figure 9).

## Discussion

In this study, we utilized taxon-specific high-resolution amplicons to elucidate a cross-kingdom network within the wheat phyllosphere microbiome at species and strain resolution. Since microbial community niche processes occur at small scales, we used detailed temporal and spatial profiling at a single field site throughout the growing season to capture these dynamics. Microbial interaction networks have been hypothesized to be strain-specific, a consensus derived from experimental studies involving a limited number of isolates[35, 36]. However, systematic field studies have been lacking. In this study, we addressed this gap by using taxon-specific high-resolution amplicons specific to the bacterial and fungal keystone taxa of the wheat phyllosphere, namely *Pseudomonas* and *Z. tritici*. Using state-of the art co-occurrence analysis with rigorous correction for environmental covariates, we inferred potential synergistic and antagonistic interactions among strains[40, 66]. We demonstrated a high degree of strain-specificity in *Pseudomonas* interactions across kingdoms. Through systematic analysis of negative interactions, we identified a strain-specific stable biocontrol candidate assembly of five *Pseudomonas* isolates and five fungal taxa that potentially suppresses the seven most dominant phyllosphere fungal pathogens. Furthermore, we identified potential biocontrol stabilizer taxa that exhibit positive interactions with these biocontrol candidates, potentially enhancing the effectiveness of field applications. Using a comprehensive *Pseudomonas* isolate collection from the same field, we confirmed the prevalence of positive and strain-specific interactions and further identified antagonistic effects from *Pseudomonas* strains recognized as biocontrol candidates in our network analysis.

Our network analysis revealed a dominance of positive interactions at both high and low taxonomic levels. While prior studies focusing on broader taxonomic levels have reported mixed patterns, many also found a predominance of positive associations[87, 88]. The accuracy of such estimates depends strongly on proper correction for shared environmental covariates. Commonly used correlation-based inference tools such as SparCC and CoNet do not remove indirect associations mediated by third taxa or environmental factors, compromising their interpretability[89]. In contrast, SPIEC-EASI, which we applied here, estimates direct associations by computing the inverse covariance matrix[66]. We further used its “sparse and low-rank” extension, which accounts for unobserved latent factors that may confound microbial interaction patterns [40]. To evaluate the robustness of the resulting networks, we performed subsampling analyses. Nevertheless, we cannot exclude the fact that the interactions identified here are influenced by unobserved factors. Importantly, co-occurrence analysis has inherent limitations: strongly asymmetric interactions (e.g., exploitation) often result in positive correlations, since the benefits for one partner can outweigh the costs to the other[43]. Symmetric interactions such as mutualism or competition are more likely to be correctly identified, yet even competition may appear as positive correlations when competing species differ markedly in strength[43]. This may explain why empirical networks often report relatively few negative correlations[43, 90].

Moreover, the inferred balance of positive versus negative interactions can depend on the sampling scale. Simulation studies indicate that competitive interactions become harder to detect with increasing spatial extent, whereas mutualistic, commensal, and parasitic interactions are more reliably detected across scales[91]. To mitigate these biases, and also to reduce effects of habitat filtering, sampling for biotic interactions should be derived from similar environments[39, 43]. Therefore, we chose a fine-scale spatial and temporal sampling of densely sampled plants within a single season to detect competitive interactions on finer scales. We also included different canopy heights and cultivars to encompass enough variation to detect differential co-occurrence. Nonetheless, we cannot exclude the possibility that the sampling scale skewed the actual balance of positive and negative interactions. Hence, to assess the frequency of positive and negative interactions in an orthogonal approach, we performed co-inoculation experiments of *Pseudomonas* and *Z. tritici in planta* and measured the biomass change of individual strains upon co-inoculation using qPCR. Consistent with the predictions from the field, we found that the majority of interactions were positive, that is increased biomass upon co-inoculation. It has been suggested that a low fraction of negative interactions compared to positive interactions is associated with heterogeneous microenvironments that reduce direct competition and promote niche sharing[88]. Simulations have suggested that mutualism can be maintained under infection by multiple symbionts when shared costs are sufficiently low, while greater virulence and parasitism toward the host are more likely when shared costs are high[92]. Furthermore, both positive and negative interactions pose additional evolutionary constraints and hence influence the adaptive trajectories of individual species[93]. Across all taxonomic levels, we found interaction networks to have a scale-free topology, that is a few hub nodes with many interactions and a large number of nodes with only a few interactions. This organization is most commonly found in biological and biomedical networks and has been proposed to promote robustness of microbial ecosystems[94]. Consistent with this, we found overall relative taxa abundances to be stable over timepoints, cultivars and canopy heights. An overall consistent taxonomic composition of the plant associated bacterial community was previously observed and has been attributed to a deterministic assembly[95–98]. Here we identified *Pseudomonas* and *Zymoseptoria* as hub nodes in the bacterial-fungal interaction network with an especially large number of interactions. Consequently, we aimed to investigate these interactions more closely at a higher taxonomic resolution.

We systematically tested for potential biocontrol assemblies among *Pseudomonas* species. Unlike controlled experimental co-inoculation studies, we assessed interactions at the strain level within a complex field system. Despite the prevalence of positive interactions, we identified an assembly of ten taxa that potentially suppress seven fungal pathogens, including three documented wheat pathogens: *Zymoseptoria*[21], *Blumeria*[99] and *Neoascochyta*[100]^99^. The ten suppressor taxa comprised five *Pseudomonas* species and five fungal genera. Among the *Pseudomonas* ASVs, three belonged to the *P. fluorescens* group, known for its biocontrol activity against fungal pests[27–29, 31, 101–105]. We identified a *P. poae* strain showing a negative interaction with *Zymoseptoria* and a *P. poae* and a *P. chlororaphis subsp. piscium* strain showing negative interactions with *Cladosporium*. In our co-inoculation experiments, a *P. poae* isolate also exhibited the highest antifungal activity against *Z. tritici*. This indicates that although interactions are strain-specific, there might be a prevalence within groups. However, a more extensive experimental assessment would be required to evaluate the prevalence of negative interactions among *P. poae* isolates compared to other species. Although *P. poae* strains have not been characterized for their biocontrol activity against *Z. tritici*, the *P. poae* strain CO has shown strong antifungal activity *in vitro* and under greenhouse conditions against *F. graminearum* strain PH-1 on wheat[105]. The mechanisms of protection against *F. graminearum* are not yet known, but the *P. poae* strain CO was capable of producing amylases, lipases, proteases, and cellulases, likely contributing to competition against the pathogen. Other *P. poae* strains have demonstrated various biocontrol activities on maize, soybean, switchgrass, sugar beet, grapevine, and apple[106–111]. Further inoculation experiments and metabolic characterizations of the *P. poae* isolates collected here could reveal interaction mechanisms with *Z. tritici*. The other *Pseudomonas* suppressor species from the *P. fluorescens* group, *P. chlororaphis*, has been extensively characterized for its biocontrol potential against several fungal pathogens in the rhizosphere and soil, but its activity in the phyllosphere is not well understood[29, 30, 112]. However, one study found that *P. chlororaphis* strain Q3 reduced *Blumeria* incidence on wheat leaves, with the antagonism differing between wheat cultivars[113]. Here, we found that the *P. chlororaphis* suppressor strain showed considerable abundance variation between cultivars. Antagonistic interactions between *Cladosporium* and *Pseudomonas* strains have been shown for a *P. aeruginosa* and a *P. putida* strain *in vitro*[114, 115]. The five fungal suppressors in our network showed negative interactions with *Zymoseptoria*, *Blumeria*, *Cladosporium*, *Neoascochyta*, *Leveillula* and *Monographella*. Although antagonistic behavior of the fungal suppressors against species within these genera has not been described, antifungal activity against other genera has been reported for the fungal suppressors *Sporobolomyces*, *Leucosporidium*, and *Bullera*[116–118]. Specifically, a strain of the *Sporoborolymces* genus has been shown to degrade Patulin, a mycotoxin that contaminates pome fruits and derived products worldwide[118]. A *Leucosporidium* yeast species has been shown to produce both soluble and volatile antifungal compounds[117, 119]. The antifungal action of a *Bullera* yeast strain was suggested to be mediated by induced resistance of the plant[116]. These interactions could represent symmetric interference competitions, which could be detected as negative interactions by our co-occurrence analysis. Phyllosphere yeasts have diverse adaptive traits to persist in the phyllosphere, such as biofilm formation, carbon and nitrogen acquisition, and the production of pigments and killer toxins[120, 121]. More recently, they have been recognized for their biocontrol potential[120–122]. In sum, even with the inherent challenges and possible biases in co-occurrence modelling, our analysis was able to pinpoint numerous plausible biocontrol candidates that may suppress pathogens in a stand-alone analysis.

In addition to our bacterial and fungal pathogen suppressor strains, we identified both synergistic and antagonistic interactions that potentially promote their stability or conversely inhibit the suppressor strains. A main limitation of the application of biocontrol assemblies in the field has been the lack of stable colonization[123]. It has been proposed that multi-strain biocontrol inoculants could improve niche overlap and stable establishment[123, 124]. When designing multi-strain assemblies, the compatibility of interaction mechanisms has been shown to be critical. A powerful approach could be to identify mutualistic strains that improve each other’s colonization[125]. For example, a study showed improved biocontrol efficiency when applying a *P. fluorescens* and a *Bacillus subtilis* strain together[126]. They found that the beneficial *Pseudomonas* strain can stimulate the *Bacillus* strain to produce surfactin through its T6SS system, which is beneficial for nutrient utilization and colonization of *Pseudomonas*. The suppressor stabilizers identified here provide good candidates for testing similar approaches, i.e. trying to construct suppressing communities where the biocontrol agents are stabilized by a community of mutualists. Alongside colonization success, another important aspect in the development of biocontrol applications has been the assessment of non-target effects[127, 128]. While conclusive non-target effect studies are challenging to conduct, several studies have indicated that biocontrol agents that suppress root pathogenic fungi generally have small-scale effects[129]. In line with this, our network showed a modular structure at all taxonomic levels, indicating that biocontrol applications likely have more localized and specific effects, contained within modules. Network analysis as conducted here could provide a useful tool to analyze non-target effects in the field.

In summary, we present the first network analysis based on high-resolution taxon-specific markers, which enabled us to construct species and strain-specific interaction networks. This allowed for a systematic search for potential pathogen suppressor candidates in the field, and in turn, their interactions with each other and with other facilitators and stabilizers. We demonstrated that the network predictions can be replicated with co-inoculation experiments with isolates from the same field, including the identification of antagonistic suppressor strains against fungal pathogens. These findings provide interesting candidates for further assessment as biocontrol agents and biocontrol consortia for agricultural applications.

## Data availability

Sequencing data generated for this work are available at the NCBI Sequence Read Archive (SRA) BioProject PRJNA1102740.

## Code availability

Analysis scripts generated for this manuscript are shared on the Github repository https://github.com/LuziaThea/wheat-microbiome-net

## Author contributions

LS, MM and DC conceived the study, LS performed the research and analyzed the data. LS and AS performed the *in vitro* and *in planta* co-inoculation experiments. MM and DC supervised the work and acquired funding. LS and DC wrote the manuscript with input from MM.

## Acknowledgments

Data produced in this paper were generated in collaboration with the Genetic Diversity Centre (GDC), ETH Zurich. We thank Silvia Kobel and Aria Minder for the helpful discussions regarding the DNA extraction and PCR automations. We thank Lina Jäger, Stojanka Mitrovic and Amélie Olsen for the help with the *in vitro* and *in planta* experiments. We thank Dominic Stalder and Sabina Tralamazza for all the helpful discussions on the analysis and Hanspeter Stalder for the input on qPCR design and troubleshooting.

## References

1. Faust K, Raes J. Microbial interactions: from networks to models. Nature Reviews Microbiology 2012 10:8 2012;10:538–550. 10.1038/nrmicro2832

2. Rodríguez-Martínez JM, Pascual A. Antimicrobial resistance in bacterial biofilms. Reviews and Research in Medical Microbiology 2006;17:65–75. 10.1097/01.REVMEDMI.0000259645.20603.63

3. Woyke T et al. Symbiosis insights through metagenomic analysis of a microbial consortium. Nature 2006 443:7114 2006;443:950–955. 10.1038/nature05192

4. Leschine SB. Cellulose degradation in anaerobic environments. Annu Rev Microbiol 1995;49:399–426. 10.1146/ANNUREV.MI.49.100195.002151

5. Hassani MA, Durán P, Hacquard S. Microbial interactions within the plant holobiont. Microbiome 2018;6:58. 10.1186/s40168-018-0445-0

6. Records AR. The type VI secretion system: a multipurpose delivery system with a phage-like machinery. Mol Plant Microbe Interact 2011;24:751–757. 10.1094/MPMI-11-10-0262

7. Tyc O et al. The Ecological Role of Volatile and Soluble Secondary Metabolites Produced by Soil Bacteria. Trends Microbiol 2017;25:280–292. 10.1016/J.TIM.2016.12.002

8. Wandersman C, Delepelaire P. Bacterial iron sources: from siderophores to hemophores. Annu Rev Microbiol 2004;58:611–647. 10.1146/ANNUREV.MICRO.58.030603.123811

9. Little AEF et al. Rules of engagement: interspecies interactions that regulate microbial communities. Annu Rev Microbiol 2008;62:375–401. 10.1146/ANNUREV.MICRO.030608.101423

10. Mercado-Blanco J, Bakker PAHM. Interactions between plants and beneficial Pseudomonas spp.: exploiting bacterial traits for crop protection. Antonie Van Leeuwenhoek 2007;92:367–389. 10.1007/S10482-007-9167-1

11. Raes J, Foerstner KU, Bork P. Get the most out of your metagenome: computational analysis of environmental sequence data. Curr Opin Microbiol 2007;10:490–498. 10.1016/J.MIB.2007.09.001

12. Blanchet FG, Cazelles K, Gravel D. Co-occurrence is not evidence of ecological interactions. Ecol Lett 2020;23:1050–1063. 10.1111/ELE.13525

13. Fones HN et al. Threats to global food security from emerging fungal and oomycete crop pathogens. Nat Food 2020;1:332–342. 10.1038/s43016-020-0075-0

14. Syed Ab Rahman SF et al. Emerging microbial biocontrol strategies for plant pathogens. Plant Science 2018;267:102–111. 10.1016/J.PLANTSCI.2017.11.012

15. Ørsted M et al. Population bottlenecks constrain host microbiome diversity and genetic variation impeding fitness. PLoS Genet 2022;18:e1010206. 10.1371/JOURNAL.PGEN.1010206

16. Eisenhauer N et al. Niche dimensionality links biodiversity and invasibility of microbial communities. Funct Ecol 2013;27:282–288. 10.1111/J.1365-2435.2012.02060.X

17. Fisher MC et al. Emerging fungal threats to animal, plant and ecosystem health. Nature 2012;484:186–194. 10.1038/nature10947

18. Kerdraon L et al. Differential dynamics of microbial community networks help identify microorganisms interacting with residue-borne pathogens: The case of Zymoseptoria tritici in wheat. Microbiome 2019;7. 10.1186/S40168-019-0736-0

19. Sapkota R, Jørgensen LN, Nicolaisen M. Spatiotemporal variation and networks in the mycobiome of the wheat canopy. Front Plant Sci 2017;8. 10.3389/fpls.2017.01357

20. Barroso-Bergadà D et al. Metagenomic Next-Generation Sequencing (mNGS) Data Reveal the Phyllosphere Microbiome of Wheat Plants Infected by the Fungal Pathogen Zymoseptoria tritici. Phytobiomes J 2023;7:281–287. 10.1094/PBIOMES-02-22-0008-FI/SUPPL_FILE/PBIOMES-02-22-0008-FI.ST1.PDF

21. Dean R et al. The Top 10 fungal pathogens in molecular plant pathology. Mol Plant Pathol 2012;13:414–430. 10.1111/J.1364-3703.2011.00783.X

22. Fones H, Gurr S. The impact of Septoria tritici Blotch disease on wheat: An EU perspective. Fungal Genet Biol 2015;79:3–7. 10.1016/J.FGB.2015.04.004

23. Legein M et al. Modes of Action of Microbial Biocontrol in the Phyllosphere. Front Microbiol 2020;11. 10.3389/fmicb.2020.01619

24. Müller T, Behrendt U. Exploiting the biocontrol potential of plant-associated pseudomonads – A step towards pesticide-free agriculture? Biological Control 2021;155:104538. 10.1016/J.BIOCONTROL.2021.104538

25. Kavamura VN et al. Defining the wheat microbiome: Towards microbiome-facilitated crop production. Comput Struct Biotechnol J 2021;19:1200. 10.1016/J.CSBJ.2021.01.045

26. Chen J, et al. Wheat Microbiome: Structure, Dynamics, and Role in Improving Performance Under Stress Environments. Front Microbiol 2021;12. 10.3389/FMICB.2021.821546

27. Haas D, Défago G. Biological control of soil-borne pathogens by fluorescent pseudomonads. Nat Rev Microbiol 2005;3:307–319. 10.1038/nrmicro1129

28. Pieterse CMJ et al. Induced systemic resistance by beneficial microbes. Annu Rev Phytopathol 2014;52:347–375. 10.1146/ANNUREV-PHYTO-082712-102340/CITE/REFWORKS

29. Anderson AJ, Kim YC. Insights into plant-beneficial traits of probiotic Pseudomonas chlororaphis isolates. J Med Microbiol 2020;69:361–371. 10.1099/JMM.0.001157

30. Arrebola E et al. Fitness features involved in the biocontrol interaction of pseudomonas chlororaphiswith host plants: The case study of PcPCL1606. Front Microbiol 2019;10:1–8. 10.3389/fmicb.2019.00719

31. Raio A, Puopolo G. Pseudomonas chlororaphis metabolites as biocontrol promoters of plant health and improved crop yield. World J Microbiol Biotechnol 2021;37. 10.1007/S11274-021-03063-W

32. Zhang Y et al. Volatile Organic Compounds Produced by Pseudomonas chlororaphis subsp. aureofaciens SPS-41 as Biological Fumigants to Control Ceratocystis fimbriata in Postharvest Sweet Potatoes. J Agric Food Chem 2019;67:3702–3710. 10.1021/acs.jafc.9b00289

33. Gislason AS, de Kievit TR. Friend or foe? Exploring the fine line between Pseudomonas brassicacearum and phytopathogens. J Med Microbiol 2020;69:347–360. 10.1099/JMM.0.001145

34. Xin XF, Kvitko B, He SY. Pseudomonas syringae: What it takes to be a pathogen. Nat Rev Microbiol 2018;16:316–328. 10.1038/NRMICRO.2018.17

35. Vishwakarma K et al. Revisiting Plant–Microbe Interactions and Microbial Consortia Application for Enhancing Sustainable Agriculture: A Review. Front Microbiol 2020;11:560406. 10.3389/FMICB.2020.560406/BIBTEX

36. Gupta R et al. Plant–microbiome interactions for sustainable agriculture: a review. Physiology and Molecular Biology of Plants 2021;27:165–179. 10.1007/S12298-021-00927-1/FIGURES/4

37. McEvoy PB. Theoretical contributions to biological control success. BioControl 2018;63:87–103. 10.1007/S10526-017-9852-6/TABLES/1

38. Schulz AN, Lucardi RD, Marsico TD. Successful Invasions and Failed Biocontrol: The Role of Antagonistic Species Interactions. Bioscience 2019;69:711–724. 10.1093/BIOSCI/BIZ075

39. Berry D, Widder S. Deciphering microbial interactions and detecting keystone species with co-occurrence networks. Front Microbiol 2014;5:90985. 10.3389/FMICB.2014.00219/BIBTEX

40. Kurtz ZD, Bonneau R, Müller CL. Disentangling microbial associations from hidden environmental and technical factors via latent graphical models. BioRxiv 2019. 10.1101/2019.12.21.885889

41. McGill BJ. Ecology. Matters of scale. Science 2010;328:575–576. 10.1126/SCIENCE.1188528

42. Russell R et al. Scale, environment, and trophic status: the context dependency of community saturation in rocky intertidal communities. Am Nat 2006;167:158–170. 10.1086/504603/ASSET/IMAGES/LARGE/FG21_ONLINE.JPEG

43. Pinto S et al. Species abundance correlations carry limited information about microbial network interactions. PLoS Comput Biol 2022;18:e1010491. 10.1371/JOURNAL.PCBI.1010491

44. Liu YX et al. A practical guide to amplicon and metagenomic analysis of microbiome data. Protein Cell 2021;12:315–330. 10.1007/S13238-020-00724-8

45. Earl JP et al. Species-level bacterial community profiling of the healthy sinonasal microbiome using Pacific Biosciences sequencing of full-length 16S rRNA genes. Microbiome 2018;6. 10.1186/S40168-018-0569-2

46. Stalder L et al. High-resolution profiling of bacterial and fungal communities using pangenome-informed taxon-specific amplicons and long-read sequencing. Microbiome 2025;in press.

47. Large EC. Growth stages in cereals - illustration of the Feekes scale. Plant Pathol 1954;3:128–129. 10.1111/J.1365-3059.1954.TB00716.X

48. Zenkl R, Anderegg J, McDonald B. leaf-toolkit: Leaf Evaluation and Analysis Framework (LEAF). https://github.com/RadekZenkl/leaf-toolkit. (2023, date last accessed).

49. Altschul SF et al. Basic local alignment search tool. J Mol Biol 1990;215:403–410. 10.1016/S0022-2836(05)80360-2

50. Martin M. Cutadapt removes adapter sequences from high-throughput sequencing reads. EMBnet J 2011;17:10–12. 10.14806/EJ.17.1.200

51. Callahan BJ et al. DADA2: High-resolution sample inference from Illumina amplicon data. Nature Methods 2016 13:7 2016;13:581–583. 10.1038/nmeth.3869

52. Yilmaz P et al. The SILVA and “All-species Living Tree Project (LTP)” taxonomic frameworks. Nucleic Acids Res 2014;42:D643–D648. 10.1093/NAR/GKT1209

53. Nilsson RH et al. The UNITE database for molecular identification of fungi: handling dark taxa and parallel taxonomic classifications. Nucleic Acids Res 2019;47:D259–D264. 10.1093/NAR/GKY1022

54. Shen W et al. SeqKit: A Cross-Platform and Ultrafast Toolkit for FASTA/Q File Manipulation. PLoS One 2016;11:e0163962. 10.1371/JOURNAL.PONE.0163962

55. Winsor GL et al. Enhanced annotations and features for comparing thousands of Pseudomonas genomes in the Pseudomonas genome database. Nucleic Acids Res 2016;44:D646–D653. 10.1093/NAR/GKV1227

56. Lalucat J et al. Genomics in bacterial taxonomy: Impact on the genus pseudomonas. Genes (Basel) 2020;11. 10.3390/genes11020139

57. Singh NK, Karisto P, Croll D. Population-level deep sequencing reveals the interplay of clonal and sexual reproduction in the fungal wheat pathogen Zymoseptoria tritici. Microb Genom 2021;7:678. 10.1099/mgen.0.000678

58. Badet T et al. A 19-isolate reference-quality global pangenome for the fungal wheat pathogen Zymoseptoria tritici. BMC Biol 2020;18:1–18. 10.1186/s12915-020-0744-3

59. Nguyen NH et al. FUNGuild: An open annotation tool for parsing fungal community datasets by ecological guild. Fungal Ecol 2016;20:241–248. 10.1016/j.funeco.2015.06.006

60. King EO, Ward MK, Raney DE. Two simple media for the demonstration of pyocyanin and fluorescin. J Lab Clin Med 1954;2:301–307.

61. Bertani G. Studies on lysogenesis. I. The mode of phage liberation by lysogenic Escherichia coli. J Bacteriol 1951;62:293–300.

62. Keel C et al. Conservation of the 2,4-diacetylphloroglucinol biosynthesis locus among fluorescent Pseudomonas strains from diverse geographic locations. Appl Environ Microbiol 1996;62:552–563. 10.1128/AEM.62.2.552-563.1996

63. Stutz EW. Naturally Occurring Fluorescent Pseudomonads Involved in Suppression of Black Root Rot of Tobacco. Phytopathology 1986;76:181. 10.1094/PHYTO-76-181

64. Flury P et al. Insect pathogenicity in plant-beneficial pseudomonads: Phylogenetic distribution and comparative genomics. ISME Journal 2016;10:2527–2542. 10.1038/ismej.2016.5

65. Chin-A-Woeng TFC et al. Introduction of the phzH gene of Pseudomonas chlororaphis PCL1391 extends the range of biocontrol ability of phenazine-1-carboxylic acid-producing Pseudomonas spp. strains. Mol Plant Microbe Interact 2001;14:1006–1015. 10.1094/MPMI.2001.14.8.1006

66. Kurtz ZD et al. Sparse and Compositionally Robust Inference of Microbial Ecological Networks. PLoS Comput Biol 2015;11. 10.1371/JOURNAL.PCBI.1004226

67. Gábor Csárdi et al. Network Analysis and Visualization in R. 2024. 2024.

68. Gabor Csardi, Tamas Nepusz. The igraph software package for complex network research. InterJournal 2006;Complex Systems:1695.

69. Shannon P et al. Cytoscape: A software Environment for integrated models of biomolecular interaction networks. Genome Res 2003;13:2498–2504. 10.1101/gr.1239303

70. Mirarab S et al. PASTA: Ultra-Large Multiple Sequence Alignment for Nucleotide and Amino-Acid Sequences. Journal of Computational Biology 2015;22:377. 10.1089/CMB.2014.0156

71. Kozlov AM et al. RAxML-NG: a fast, scalable and user-friendly tool for maximum likelihood phylogenetic inference. Bioinformatics 2019;35:4453–4455. 10.1093/BIOINFORMATICS/BTZ305

72. Price MN, Dehal PS, Arkin AP. FastTree: Computing Large Minimum Evolution Trees with Profiles instead of a Distance Matrix. Mol Biol Evol 2009;26:1641. 10.1093/MOLBEV/MSP077

73. Best DJ, Roberts DE. Algorithm AS 89: The Upper Tail Probabilities of Spearman’s Rho. Appl Stat 1975;24:377. 10.2307/2347111

74. Oksanen Jari. vegan: Community Ecology Package. 2022. 2022.

75. R Core Team. R: A Language and Environment for Statistical Computing. https://www.r-project.org/. (2022, date last accessed).

76. McMurdie PJ, Holmes S. phyloseq: An R Package for Reproducible Interactive Analysis and Graphics of Microbiome Census Data. PLoS One 2013;8:e61217. 10.1371/JOURNAL.PONE.0061217

77. Curry E, Zhang H. Data Integration, Manipulation and Visualization of Phylogenetic Trees. Data Integration, Manipulation and Visualization of Phylogenetic Trees. Chapman and Hall/CRC, 2022.

78. Yu G. Using ggtree to Visualize Data on Tree-Like Structures. Curr Protoc Bioinformatics 2020;69. 10.1002/CPBI.96

79. Wickham H. ggplot2: Elegant Graphics for Data Analysis. Cham: Springer International Publishing, 2016.

80. Baquero F et al. The Origin of Niches and Species in the Bacterial World. Front Microbiol 2021;12:657986. 10.3389/FMICB.2021.657986/BIBTEX

81. Xiong C et al. Plant developmental stage drives the differentiation in ecological role of the maize microbiome. Microbiome 2021;9:1–15. 10.1186/S40168-021-01118-6/FIGURES/5

82. Vorholt JA. Microbial life in the phyllosphere. Nat Rev Microbiol 2012;10:828–840. 10.1038/NRMICRO2910

83. Laforest-Lapointe I, Messier C, Kembel SW. Host species identity, site and time drive temperate tree phyllosphere bacterial community structure. Microbiome 2016;4:1–10. 10.1186/S40168-016-0174-1/FIGURES/4

84. Noble AS et al. A core phyllosphere microbiome exists across distant populations of a tree species indigenous to New Zealand. PLoS One 2020;15:e0237079. 10.1371/JOURNAL.PONE.0237079

85. Grilli J, Rogers T, Allesina S. Modularity and stability in ecological communities. Nat Commun 2016;7. 10.1038/NCOMMS12031

86. Li L et al. Investigating the diversity of pseudomonas spp. in soil using culture dependent and independent techniques. Curr Microbiol 2013;67:423–430. 10.1007/S00284-013-0382-X/FIGURES/3

87. Huang F, Lei M, Li W. The rhizosphere and root selections intensify fungi-bacteria interaction in abiotic stress-resistant plants. PeerJ 2024;12. 10.7717/PEERJ.17225

88. Ma B et al. Earth microbial co-occurrence network reveals interconnection pattern across microbiomes. Microbiome 2020;8:1–12. 10.1186/S40168-020-00857-2/FIGURES/6

89. Röttjers L, Faust K. From hairballs to hypotheses–biological insights from microbial networks. FEMS Microbiol Rev 2018;42:761–780. 10.1093/FEMSRE/FUY030

90. Ma Z (Sam). The P/N (Positive-to-Negative Links) Ratio in Complex Networks—A Promising In Silico Biomarker for Detecting Changes Occurring in the Human Microbiome. Microb Ecol 2018;75:1063– 1073. 10.1007/S00248-017-1079-7/TABLES/5

91. Araújo MB, Rozenfeld A. The geographic scaling of biotic interactions. Ecography 2014;37:406–415. 10.1111/J.1600-0587.2013.00643.X

92. Nelson PG, May G. Coevolution between Mutualists and Parasites in Symbiotic Communities May Lead to the Evolution of Lower Virulence. 10.5061/dryad.dd414

93. Barber JN et al. Species interactions constrain adaptation and preserve ecological stability in an experimental microbial community. 2022. 10.1038/s41396-022-01191-1

94. Luo M et al. Progress on network modeling and analysis of gut microecology: a review. Appl Environ Microbiol 2024;90. 10.1128/AEM.00092-24/ASSET/426AB89F-DA69-47AE-A59D-2665ACA17859/ASSETS/IMAGES/LARGE/AEM.00092-24.F002.JPG

95. Delmotte N et al. Community proteogenomics reveals insights into the physiology of phyllosphere bacteria. Proc Natl Acad Sci U S A 2009;106:16428. 10.1073/PNAS.0905240106

96. Wang J et al. Global assembly of microbial communities. mSystems 2023;8. 10.1128/MSYSTEMS.01289-22/SUPPL_FILE/MSYSTEMS.01289-22-S0010.XLSX

97. Huang S, Zha X, Fu G. Affecting Factors of Plant Phyllosphere Microbial Community and Their Responses to Climatic Warming—A Review. Plants 2023, Vol 12, Page 2891 2023;12:2891. 10.3390/PLANTS12162891

98. Bechtold EK et al. Successional changes in bacterial phyllosphere communities are plant-host species dependent. Appl Environ Microbiol 2024;90. 10.1128/AEM.01750-23

99. Basandrai AK, Mehta A, Basandrai D. Virulence structure of wheat powdery mildew pathogen, Blumeria graminis tritici: a review. Indian Phytopathology 2022 76:1 2022;76:21–45. 10.1007/S42360-022-00571-Z

100. Golzar H et al. Neoascochyta species cause leaf scorch on wheat in Australia. Australas Plant Dis Notes 2019;14:1–5. 10.1007/S13314-018-0332-3/FIGURES/4

101. Chin-A-Woeng TFC, Bloemberg G V., Lugtenberg BJJ. Phenazines and their role in biocontrol by Pseudomonas bacteria. New Phytologist 2003;157:503–523. 10.1046/J.1469-8137.2003.00686.X

102. Weller DM. Pseudomonas biocontrol agents of soilborne pathogens: looking back over 30 years. Phytopathology 2007;97:250–256. 10.1094/PHYTO-97-2-0250

103. Sitaraman R. Pseudomonas spp. as models for plant-microbe interactions. Front Plant Sci 2015;6:1–4. 10.3389/fpls.2015.00787

104. Biessy A, Filion M. Phenazines in plant-beneficial Pseudomonas spp.: biosynthesis, regulation, function and genomics. Environ Microbiol 2018;20:3905–3917. 10.1111/1462-2920.14395

105. Ibrahim E et al. Biocontrol Efficacy of Endophyte Pseudomonas poae to Alleviate Fusarium Seedling Blight by Refining the Morpho-Physiological Attributes of Wheat. Plants 2023, Vol 12, Page 2277 2023;12:2277. 10.3390/PLANTS12122277

106. Niem JM et al. Biocontrol Potential of an Endophytic Pseudomonas poae Strain against the Grapevine Trunk Disease Pathogen Neofusicoccum luteum and Its Mechanism of Action. Plants (Basel) 2023;12. 10.3390/PLANTS12112132

107. Xia Y et al. Improved Draft Genome Sequence of Pseudomonas poae A2-S9, a Strain with Plant Growth-Promoting Activity. Microbiol Resour Announc 2019;8:275–294. 10.1128/MRA.00275-19

108. Zachow C et al. The Novel Lipopeptide Poaeamide of the Endophyte Pseudomonas poae RE*1-1-14 Is Involved in Pathogen Suppression and Root Colonization. Mol Plant Microbe Interact 2015;28:800–810. 10.1094/MPMI-12-14-0406-R

109. Yin C et al. Wheat Rhizosphere-Derived Bacteria Protect Soybean from Soilborne Diseases. 101094/PDIS-08-23-1713-RE 2024. 10.1094/PDIS-08-23-1713-RE

110. Ren Y et al. Isolation and characterization of a Pseudomonas poae JSU-Y1 with patulin degradation ability and biocontrol potential against Penicillium expansum. Toxicon 2021;195:1–6. 10.1016/J.TOXICON.2021.02.014

111. Hernández-León R, González-Rodríguez A, Tapia-Torres Y. Phosphorus Recycling, Biocontrol, and Growth Promotion Capabilities of Soil Bacterial Isolates from Mexican Oak Forests: An Alternative to Reduce the Use of Agrochemicals in Maize Cultivation. Applied Microbiology 2022, Vol 2, Pages 965–980 2022;2:965–980. 10.3390/APPLMICROBIOL2040074

112. Raio A, Puopolo G. Pseudomonas chlororaphis metabolites as biocontrol promoters of plant health and improved crop yield. World J Microbiol Biotechnol 2021;37:1–8. 10.1007/s11274-021-03063-w

113. Pivic Radmila et al. Bacterial antagonists Bacillus sp. Q3 and Pseudomonas chlororaphis Q16 capable to control wheat powdery mildew in wheat. Rom Biotechnol Lett 2015;20:10448–10460.

114. Li Y et al. The Genus Cladosporium: A Prospective Producer of Natural Products. International Journal of Molecular Sciences 2024, Vol 25, *Page* 1652 2024;25:1652. 10.3390/IJMS25031652

115. Wang M et al. Cladodionen Is a Potential Quorum Sensing Inhibitor Against Pseudomonas aeruginosa. Marine Drugs 2020, Vol 18, *Page* 205 2020;18:205. 10.3390/MD18040205

116. de Tenório DA et al. Biological control of Rhizoctonia solani in cowpea plants using yeast. Trop Plant Pathol 2019;44:113–119. 10.1007/S40858-019-00275-2/METRICS

117. Perez MF et al. Native Killer Yeasts as Biocontrol Agents of Postharvest Fungal Diseases in Lemons. PLoS One 2016;11:e0165590. 10.1371/JOURNAL.PONE.0165590

118. Ianiri G et al. Searching for Genes Responsible for Patulin Degradation in a Biocontrol Yeast Provides Insight into the Basis for Resistance to This Mycotoxin. Appl Environ Microbiol 2013;79:3101–3115. 10.1128/AEM.03851-12

119. Vero S et al. Evaluation of yeasts obtained from Antarctic soil samples as biocontrol agents for the management of postharvest diseases of apple (Malus × domestica). FEMS Yeast Res 2013;13:189–199. 10.1111/1567-1364.12021

120. Gouka L, Raaijmakers JM, Cordovez V. Ecology and functional potential of phyllosphere yeasts. Trends Plant Sci 2022;27:1109–1123. 10.1016/J.TPLANTS.2022.06.007

121. Gouka L, et al. Genetic, Phenotypic and Metabolic Diversity of Yeasts From Wheat Flag Leaves. Front Plant Sci 2022;13:908628. 10.3389/FPLS.2022.908628/FULL

122. Freimoser FM, et al. Biocontrol yeasts: mechanisms and applications. World J Microbiol Biotechnol 2019;35. 10.1007/S11274-019-2728-4

123. Niu B et al. Microbial Interactions Within Multiple-Strain Biological Control Agents Impact Soil-Borne Plant Disease. Front Microbiol 2020;11:585404. 10.3389/FMICB.2020.585404/BIBTEX

124. Gómez-Lama Cabanás C, Mercado-Blanco J. What Determines Successful Colonization and Expression of Biocontrol Traits at the Belowground Level? Progress in Biological Control 2020;21:31–46. 10.1007/978-3-030-53238-3_3

125. Song C, Jin K, Raaijmakers JM. Designing a home for beneficial plant microbiomes. Curr Opin Plant Biol 2021;62:102025. 10.1016/J.PBI.2021.102025

126. Jia Y et al. Synergistic biocontrol of Bacillus subtilis and Pseudomonas fluorescens against early blight disease in tomato. Appl Microbiol Biotechnol 2023;107:6071–6083. 10.1007/S00253-023-12642-W/FIGURES/6

127. Howarth FG. Non-target Effects of Biological Control Agents. Biological Control: Measures of Success 2000;369–403. 10.1007/978-94-011-4014-0_13

128. Amichot M et al. Natural products for biocontrol: review of their fate in the environment and impacts on biodiversity. Environmental Science and Pollution Research 2024 2024;1–36. 10.1007/S11356-024-33256-3

129. Winding A, Binnerup SJ, Pritchard H. Non-target effects of bacterial biological control agents suppressing root pathogenic fungi. FEMS Microbiol Ecol 2004;47:129–141. 10.1016/S0168-6496(03)00261-7

