## Supplementary Figure for "High-resolution microbial network analysis defines biocontrol consortia in the wheat phyllosphere"

Stalder *et al.*

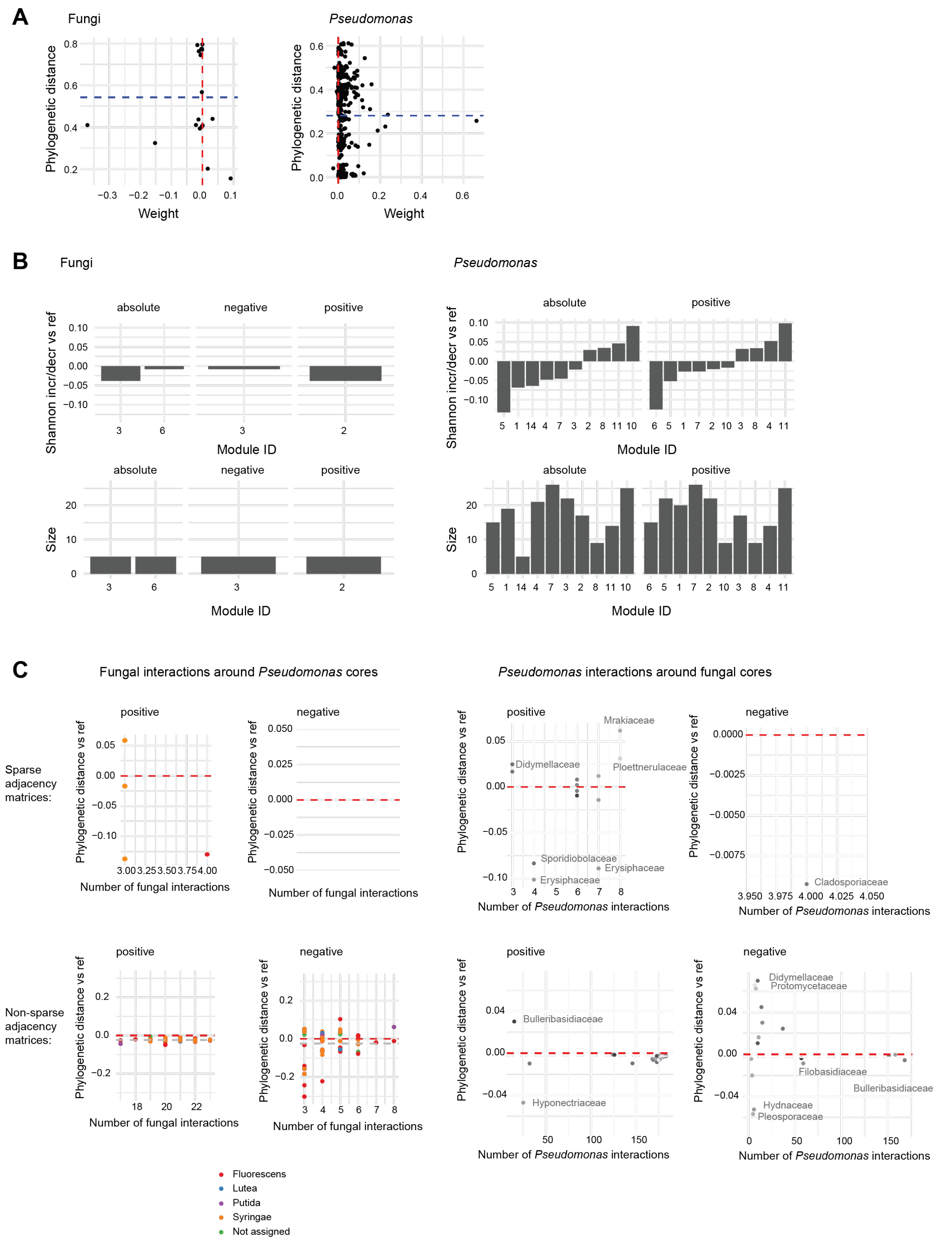


**Supplementary Figure 1**. Phylogenetic diversity of interactions in the *Pseudomonas*-fungal network. (A) Correlation between the phylogenetic distance and the interaction weight, separately for fungi and *Pseudomonas*. (B) Change in Shannon diversity of the phylogenetic diversity within network interaction modules, compared to a null expectation derived from subsampling. The phylogenetic diversity is evaluated separately for fungi (left) and *Pseudomonas* (right). The interactions within the network are considered either in absolute terms, or separated into negative and positive interactions. The size of each module is indicated at the bottom. (C) Left: Phylogenetic distances of fungal interactions shared by a single *Pseudomonas* ASV, compared to the null expectation derived from subsampling with replacement. Right: Phylogenetic distances of *Pseudomonas* interactions shared by a single fungal taxon, compared to the null expectation based on subsampling. The top row shows calculations using non-sparse interaction matrices, while the bottom row displays calculations using sparse interaction matrices.

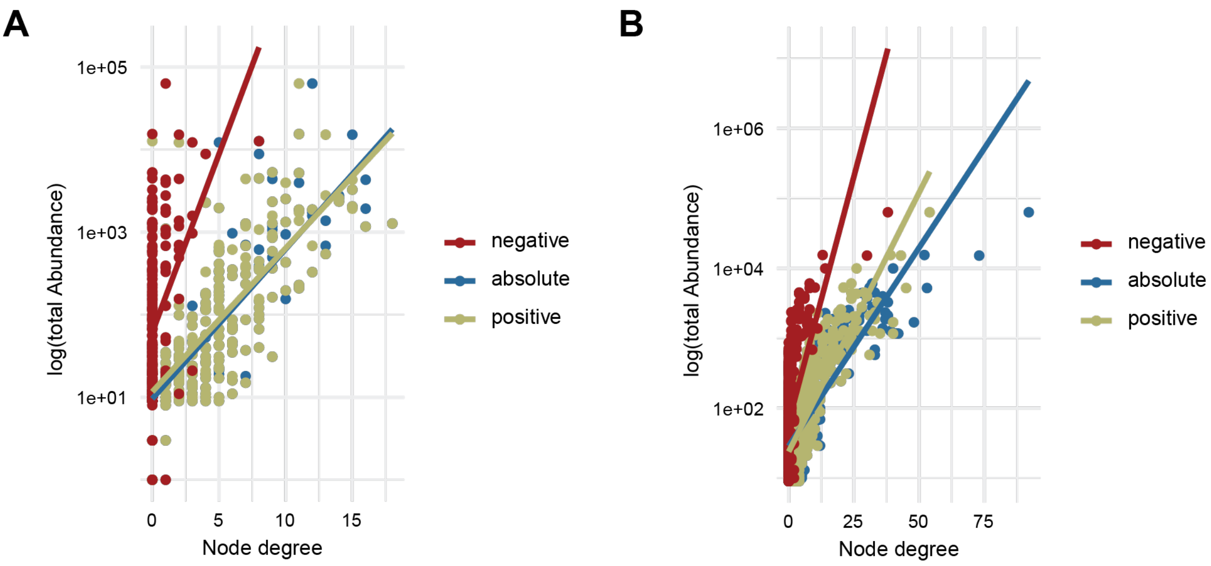


**Supplementary Figure 2**. Correlation of the general relative abundance and the node degree for the *Pseudomonas*-fungal network (A) and the *Pseudomonas-Z. tritici* network (B).

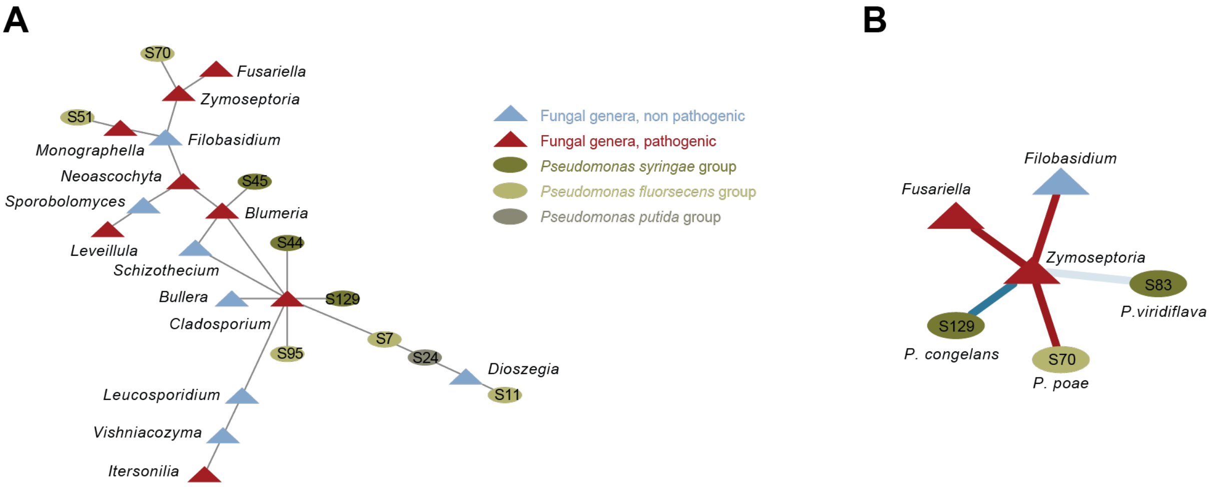


**Supplementary Figure 3**. (A) All negative interactions of the *Pseudomonas*-fungal network. (B) All direct interactions of *Zymoseptoria*.

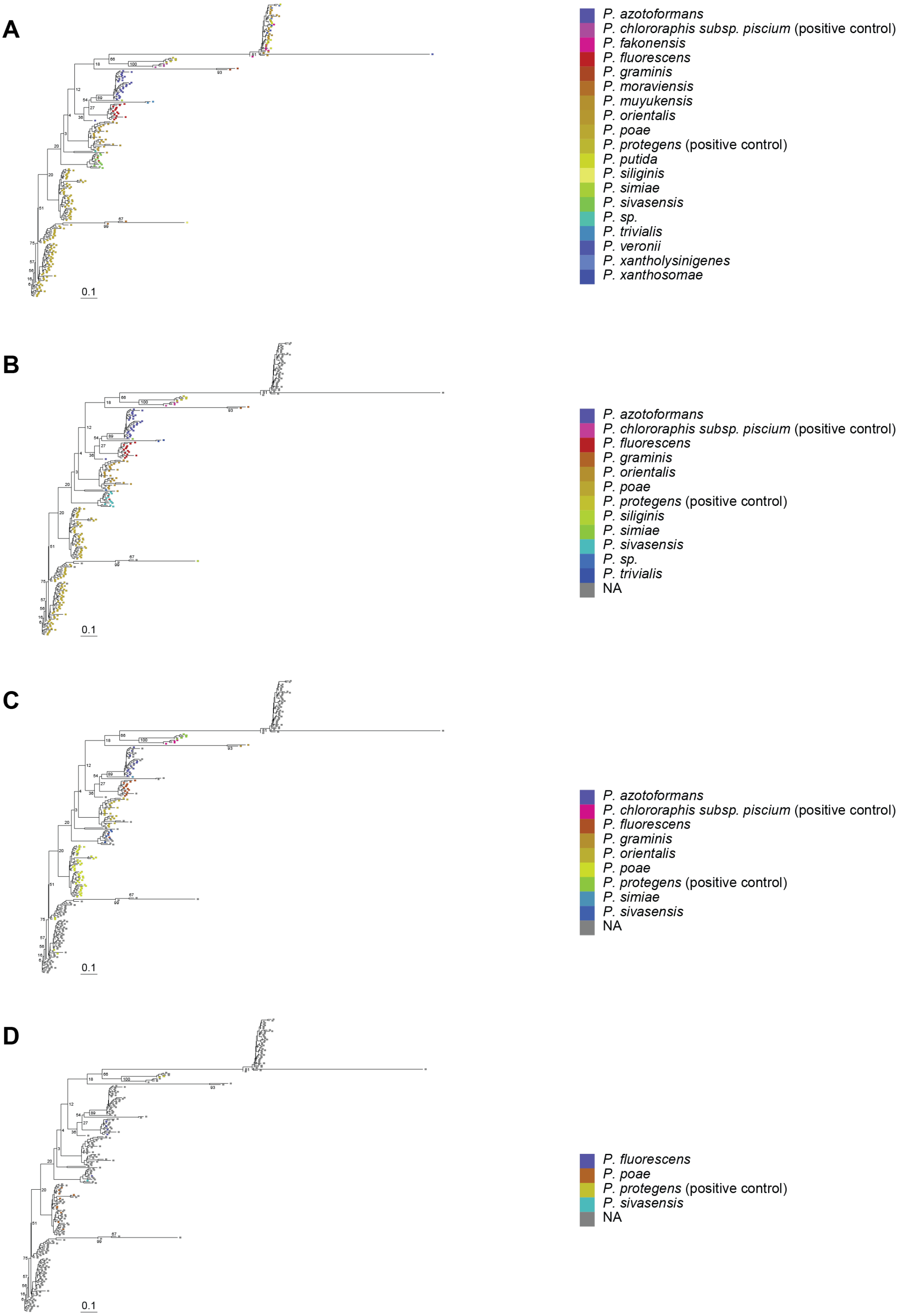


**Supplementary Figure 4**. Phylogenetic trees based on Sanger amplicon sequences of the isolates. (A) Sequences were assigned to species according to the highest BLAST bitscore. (B) Phylogenetic tree indicating only assignments with >98% identity. (C) Phylogenetic tree indicating only assignments with >95% identity. (D) Phylogenetic tree indicating only assignments with >90% identity.

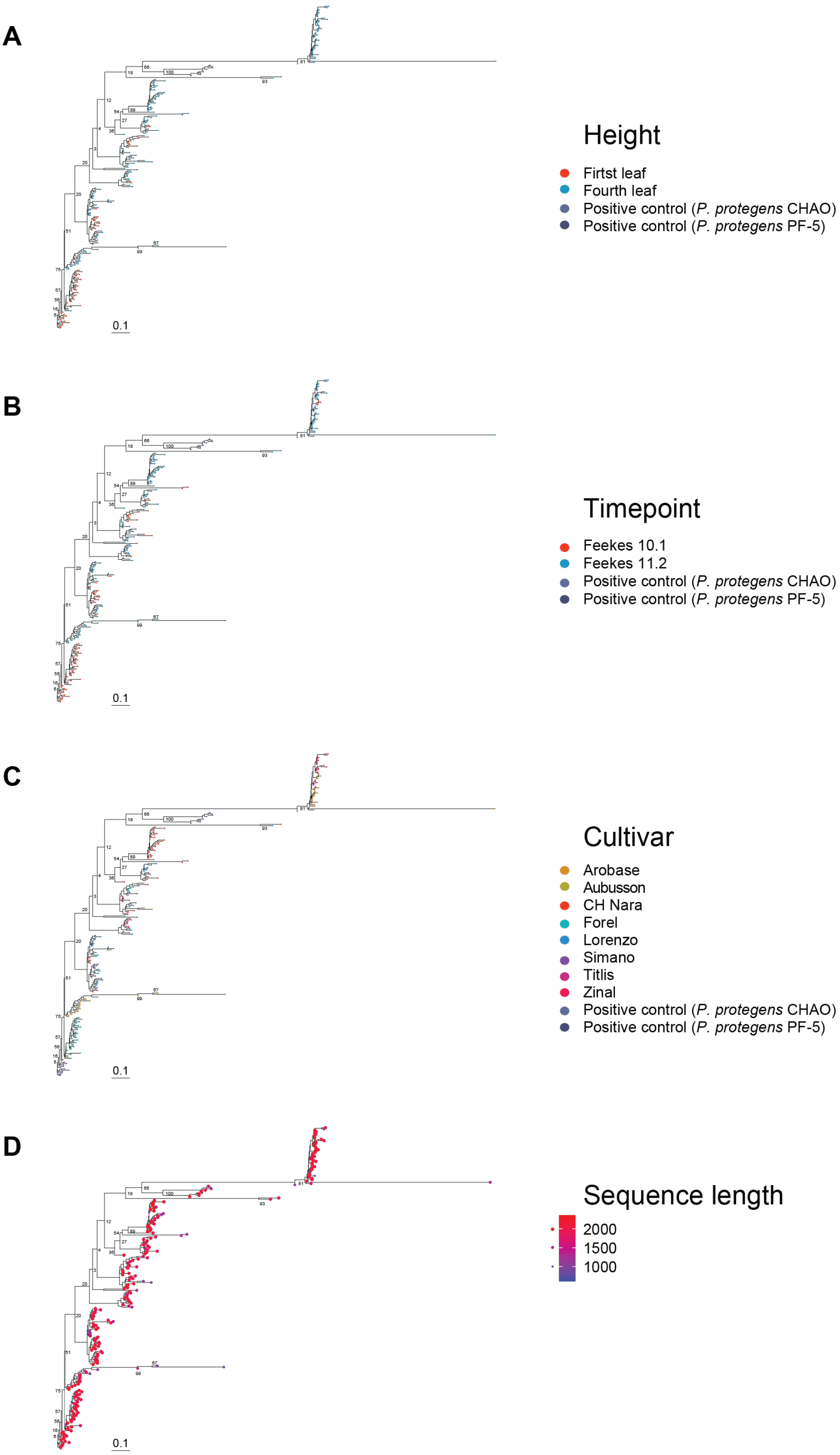


**Supplementary Figure 5**. Phylogenetic tree of Sanger amplicon sequences of the isolates. (A) Sequences are color-coded according to the canopy height from which the isolates were sampled. (B) Sequences are color-coded according to the sampling timepoint. (C) Sequences are color-coded according to the cultivar from which the isolates were obtained. (D) The length of the Sanger sequences for each isolate is indicated.

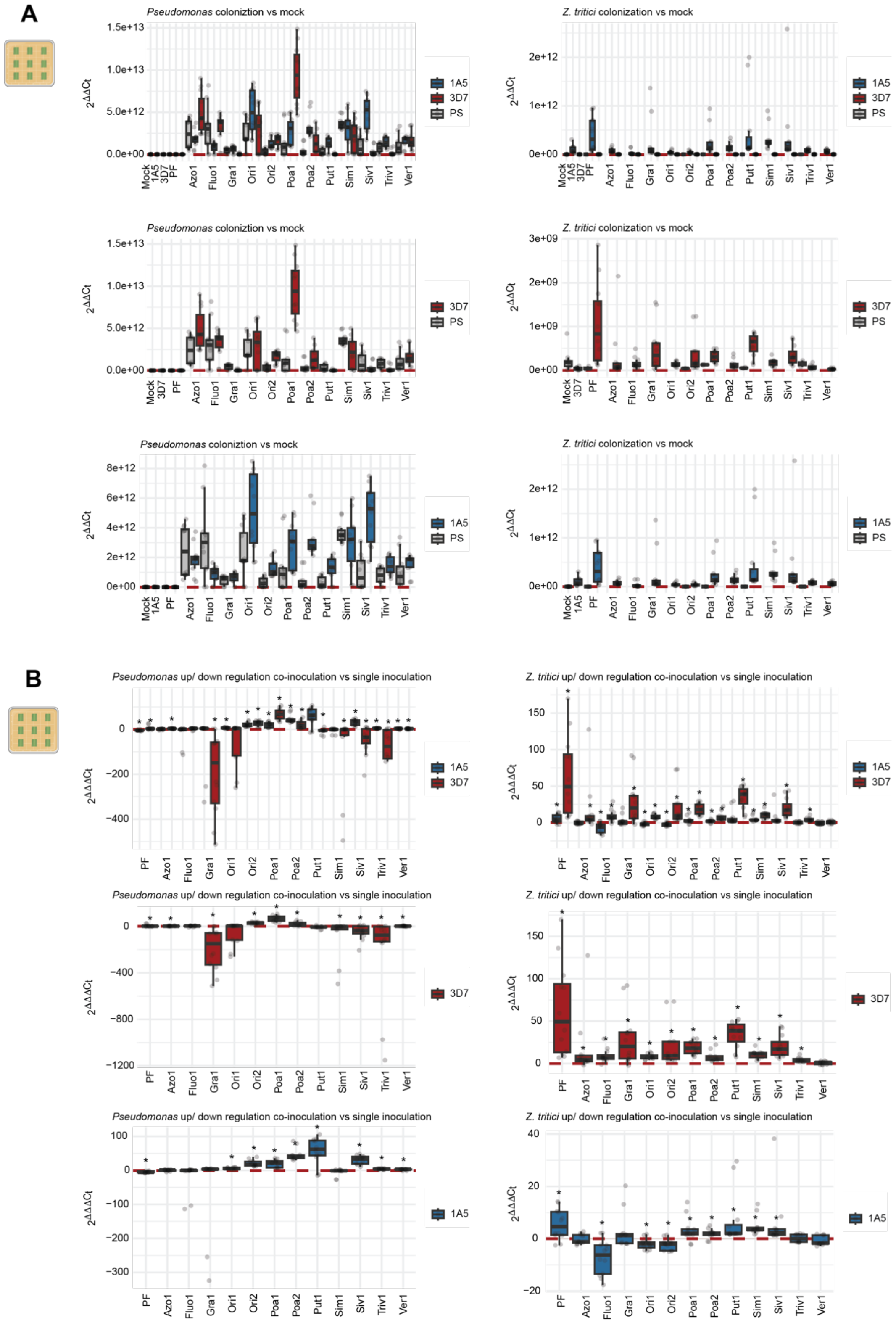


**Supplementary Figure 6**. *Pseudomonas* and *Z. tritici* biomass levels following co-inoculation with *Z. tritici* strains 1A5 and 3D7, as measured by qPCR. Isolate characteristics are detailed in Supplementary Table 9. (A) Biomass was initially normalized to wheat biomass levels and subsequently normalized relative to the mock-inoculated control, expressed as ΔΔCT. The biomass of *Z. tritici* isolate 3D7 appears 1000-fold lower compared to isolate 1A5, likely due to reduced PCR efficiency for the former. (B) Fold change of co-inoculation versus single inoculation, represented as ΔΔΔCT. Wilcoxon Rank Sum tests were conducted to determine significant differences between co-inoculations and single inoculations (p-values < 0.05).

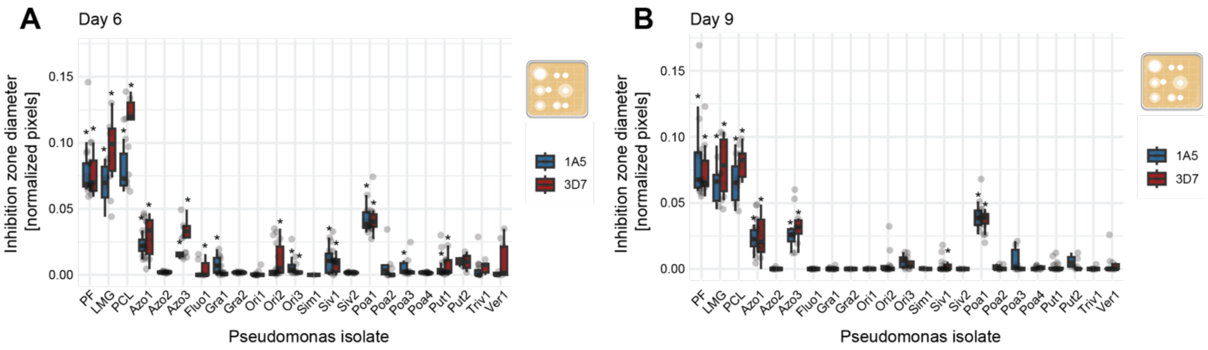


**Supplementary Figure 7**. *In vitro* growth inhibition assay demonstrating the ability of *Pseudomonas* isolates to inhibit *Z. tritici* strains 1A5 and 3D7, measured at day 5 (A) and day 9 (B). Isolate characteristics are detailed in Supplementary Table 9. Wilcoxon Rank Sum tests were conducted to determine significant differences between co-inoculations and single inoculations (p-values < 0.05).

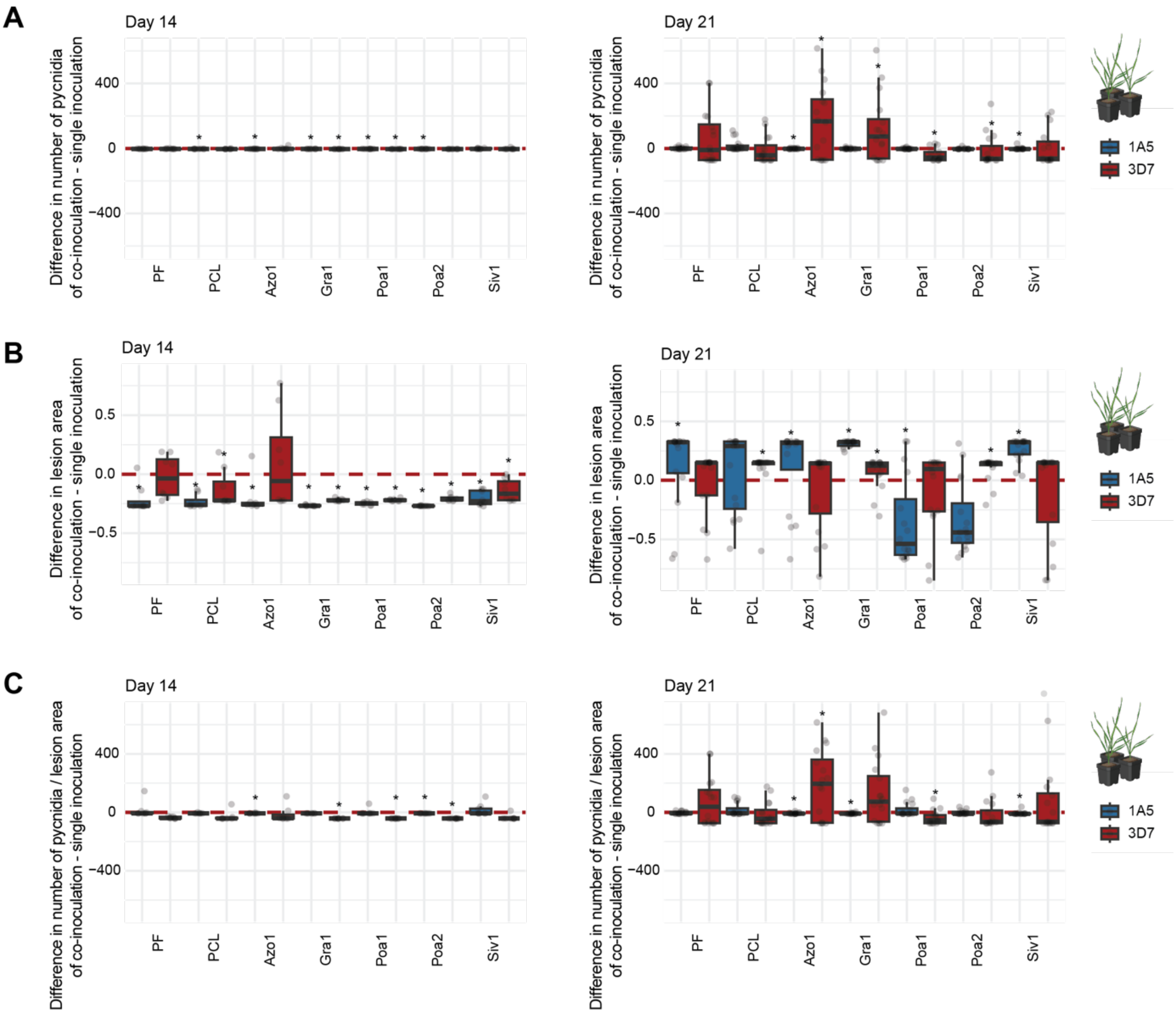


**Supplementary Figure 8**: Impact of *Pseudomonas* isolates on the disease severity of *Z. tritici* 14 and 21 days after co-inoculation. The difference of disease severity of co-inoculation – single inoculation is displayed. Measured is the number of pycnidia (A), the lesion aera (B) and the number of pycnidia per lesion area (C). Isolate characteristics are detailed in Supplementary Table 9. Wilcoxon Rank Sum tests were conducted to determine significant differences between co-inoculations and single inoculations (p-values < 0.05).

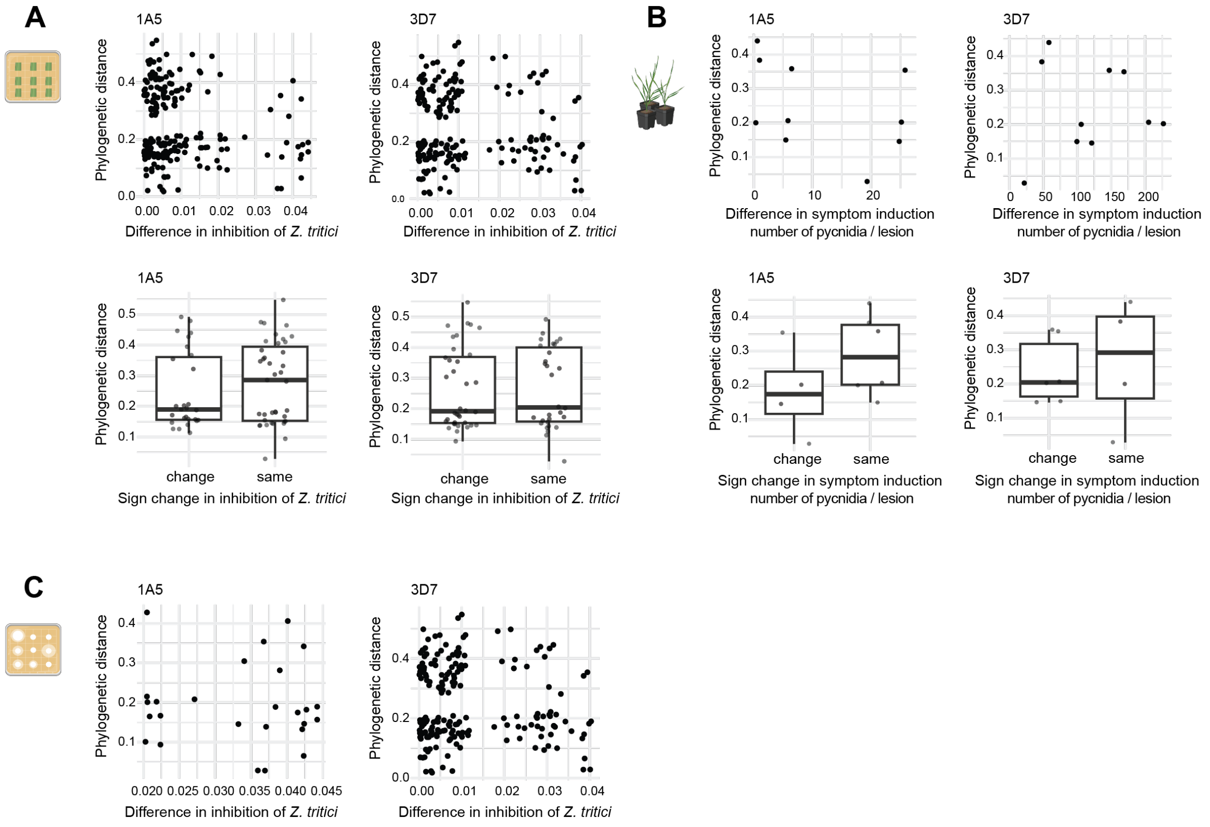


**Supplementary Figure 9**: (A) Phylogenetic distance of *Pseudomonas* isolate pairs based on their amplicon sequence compared to the difference in biomass fold change after co-inoculation with *Z. tritici* 1A5 (left) and 3D7 (right) measured by qPCR. The variation in biomass fold change is further categorized based on whether the change occurs in the same or opposite direction among pairs. (B) Phylogenetic distance of *Pseudomonas* isolate pairs based on their amplicon sequence compared to the difference in symptom induction after co-inoculation with *Z. tritici* 1A5 (left) and 3D7 (right). The variation in symptom induction is further categorized based on whether the change occurs in the same or opposite direction among pairs. (C) Phylogenetic distance of *Pseudomonas* isolate pairs based on their amplicon sequence compared to the difference in inhibition in the *in vitro* growth assay against *Z. tritici* 1A5 (left) and 3D7 (right).
